# Intraguild predation, weather, and climate teleconnection patterns interact to determine an insect vital rate

**DOI:** 10.64898/2026.08.21.746272

**Authors:** Laurent Duverglas, Carol L. Boggs

## Abstract

Population dynamics and their component vital rates may be driven by weather, climate teleconnections between sea and air (e.g. ENSO), or biotic interactions. These drivers operate directly or indirectly and on different temporal scales. We used a Bayesian structural equation model to characterize effects among weather, climate, and incidental intraguild predation (IGP) on the butterfly *Euphydryas gillettii’*s vital rate of pre-diapause survival, using an 18 year dataset. IGP was a major determinant of pre-diapause survival, along with direct and indirect effects of weather and spring climate teleconnections. The direction of climate effects was reversed when mediated through IGP. Our analysis illustrates the need for sequential hypotheses to capture the cascading effects of abiotic factors via biotic interactions. Using sequential hypotheses addresses the debate on weather – climate teleconnection roles by disentangling their contributions from one another. Finally, vital rates must be decomposed to component rates in order to detect their drivers.

## Introduction

Disentangling the contributions of biotic and abiotic factors to population dynamics is an old, yet major, goal in ecology. Insight into the relative importance of these factors can be traced back to a debate on whether populations are driven by environmental conditions (Andrewartha and Birch, 1954) or biotic mechanisms (Elton, 1949; Nicholson, 1954). Ecologists today agree that demographic processes are often shaped by a variety of biotic and abiotic factors (e.g., Price et al. 1980; Hunter et al. 1997). However, the interactions among biotic and abiotic factors are seldom tested in demographic analyses (but see Govindarajulu & Anholt, 2006; Boggs & Inouye, 2012). For instance, weather can have indirect effects on populations that are mediated through biotic interactions (White, 2008). Such indirect effects are not as readily detectable as direct effects because they require mechanistic hypotheses based on species’ biology (e.g. Benton et al. 2006; Boggs & Inouye, 2012). We document the importance of such hypotheses by showing that abiotic factors can have complex effects on vital rates (i.e., survival, growth, fecundity) via biotic interactions. Our study addresses this old debate by documenting abiotic factors whose effects on a vital rate became apparent only through their effects on biotic interactions.

Among abiotic factors, local weather conditions can be important drivers of change in vital rates (e.g. Kingsolver, 1989; van de Pol et al. 2016). Recent work has included the role of the temporal scale of weather drivers of vital rates (van de Pol et al. 2016; Hindle et al. 2019). For instance, the exact timescale on which weather is examined (e.g. monthly, annual) may reflect either acute or cumulative effects on demographic processes (e.g. Boggs and Murphy, 1997). For vital rates, it is unclear whether the true strength (i.e. overall magnitude) and/or importance (i.e. relative effect) of weather is obscured when examined over only one of these timescales. Thus, explicit hypotheses that examine weather effects occurring over a variety of timescales are necessary to identify biologically relevant weather signals that, for instance, may only show up via time lags (van de pol et al. 2016).

Temporal variability in weather can be caused by recurring transient oceanic-atmospheric anomalies that span broad geographic distributions and cause regional climatic variation over seasonal, annual, and decadal timescales (e.g. Stenseth et al. 2003; Wan et al. 2022), including the North Atlantic Oscillation (NAO), El Nino Southern Oscillation (ENSO) and many others. These climate teleconnection patterns can indirectly drive population dynamics by altering local weather conditions (e.g. Coulson et al. 2001; Stenseth et al. 2003; Roland & Matter, 2013). Teleconnection patterns can also affect populations by driving changes in biotic interactions and/or the relative relationship among the weather variables (reviewed in Wan et al. 2022). Thus, detecting the signal of teleconnection patterns on population dynamics requires identifying an underlying mechanism that connects changes in local weather conditions to ecological processes (Ottersen et al. 2001). For instance, the population dynamics of 29 North Atlantic seabird species were explained by lagged effects of the winter North Atlantic Oscillation index on adult survival and fecundity, which were respectively driven by changes in weather and fish availability (Sandvik et al. 2012; Sandvik & Erikstad, 2008). Finally, there is conflicting evidence as to whether teleconnections are better at forecasting population change than local weather conditions (Hallett et al. 2004; Frederiksen et al. 2004; Hone & Clutton-Brock, 2007; Hindle et al. 2019; Laczi et al. 2019). Our study shows that disentangling the contributions of weather from teleconnections clarifies this conflict and supports a temporally explicit understanding of demographic variation.

While biotic factors (e.g. food availability, predation, competition) alone can be key determinants of vital rates, their effects may be altered by weather (e.g. White, 2008; Mabille et al. 2010; Goodsman et al. 2018). For instance, the population dynamics of the butterfly *Speyeria mormonia* were driven by snowmelt timing which indirectly affected adult fecundity by decreasing per-capita nectar availability (Boggs & Inouye, 2012). Revealing such indirect effects of weather requires sequential hypotheses (Yang, 2020) of how demographic drivers co-vary across time. Yet sequential hypotheses remain underused in population ecology because they require long-term data (Marini et al. 2013; but see Boggs and Inouye, 2012; Juhasz et al. 2020; Duncan et al. 2021). Separating the direct effects of weather and climate from indirect effects mediated through biotic interactions will clarify the relationship between biotic and abiotic drivers of demographic change (Fig. 1).

**Figure 1.**
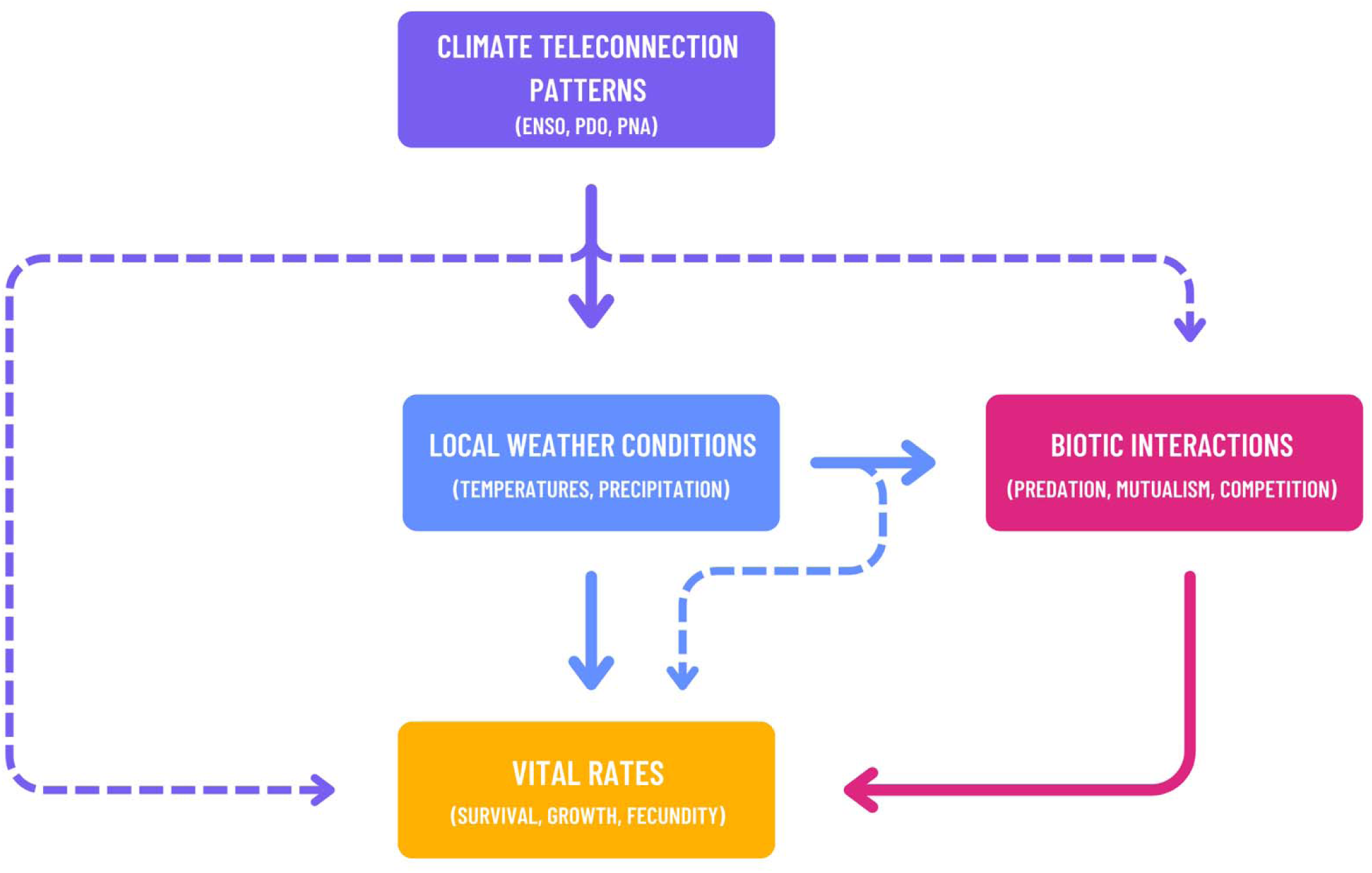
Local weather conditions and biotic interactions drive variation in vital rates through direct effects (solid arrows). Weather can also drive vital rates through indirect effects (dashed arrows) mediated via biotic interactions. Through altering local weather conditions, broadscale climate patterns indirectly drive vital rates and biotic interactions.

Among biotic interactions, intraguild predation is widespread and occurs between competitors that eat one another (Polis et al. 1989, Holt & Polis, 1997). Theoretical predictions and empirical work show that such predation can significantly alter intraguild prey population dynamics (e.g. Polis et al. 1989; Rosenheim et al. 1993; Finke & Denno, 2003). Intraguild predation is typically driven by asymmetries in size and/or age (Polis et al. 1989; Gish et al. 2017) but may also be affected by habitat complexity (Finke & Denno, 2002) and resource availability (LoPresti et al. 2018). A key, yet understudied, interaction in terrestrial communities occurs when ungulates inadvertently consume insects while foraging (Gish et al. 2017), resulting in an example of incidental intraguild predation. Short-term experiments focused on changes in abundance comprise the few studies that document ungulate-driven intraguild predation (e.g. Zamora & Gómez, 1993; Gómez & González-Megías, 2002; Bonal & Muñoz, 2007). Unlike other forms of intraguild predation, factors that drive the intensity of incidental intraguild predation apart from body size differences are relatively unknown (Polis et al. 1989; Gish et al. 2017).

Here, we evaluate the direct effects of incidental intraguild predation, weather, and climate, and the indirect effects of weather and climate via intraguild predation on an insect vital rate. We use the butterfly *Euphydryas gillettii* (Lepidoptera: Nymphalidae) and its ungulate intraguild predators. Using Bayesian structural equation modeling, we address two primary questions: (1) what are the direct and indirect effects of intraguild predation, weather, and climate on the butterfly’s vital rate of pre-diapause larval survival and (2) which among biotic and abiotic factors alter the intensity of IGP across time?

## Materials & Methods

### Study system

*Euphydryas gillettii* (Lepidoptera: Nymphalidae) inhabits the northern Rocky Mountains of North America (Williams, 1984). Our study population was introduced south of the species’ native range, to a 1.75- hectare patch in Gunnison County, Colorado at the Rocky Mountain Biological Laboratory [38.9585 N, 106.9877 W, 2,896 m a.s.l.] (Holdren & Ehrlich, 1981). The adult flight period lasts 2 - 5 weeks, typically beginning in July. Following mating, females lay eggs in clusters on the preferred host plant, *Lonicera involucrata* (Dipscales: Caprifoliaceae) (Williams et al. 1984). Egg clusters hatch after 18 – 30 days, depending on temperature and sunlight exposure (Bonebrake et al. 2010). Caterpillars feed in communal webs on *L. involucrata* prior to entering over-winter diapause a few weeks after hatching. Deer, *Odocoileus hemionus,* and elk, *Cervus canadensis,* (Artiodactyla: Cervidae) may incidentally consume egg clusters and the occasional first-instar larval web while browsing *L. involucrata*. After terminating diapause the following May, larvae feed individually before pupating to repeat the cycle (Williams et al. 1984).

### Egg cluster data and the pre-diapause survival probability

We monitored E. *gillettii* egg clusters (n ≥ 60) each year between 2006-2024. In years when it was not feasible to monitor all egg clusters due to their large number, we arbitrarily chose clusters distributed across the study area. Each cluster was marked, photographed, and the number of eggs was counted. We recorded the fate of larval webs for each marked egg cluster in early September, just prior to larval diapause. If a larval web was present, we counted the number of larvae within it. The pre-diapause survival probability was the product of the proportion of surviving larval webs and the mean proportion of surviving larvae (Morris et al. 2008). We analyzed these two components of pre-diapause survival separately because they represent different processes which may have separate drivers. We excluded data from 2012 due to an unknown predator or fungus that attacked the leaves within larval webs, turning them to mush. Based on the adult population size in the subsequent year, larvae from affected webs apparently escaped by falling to the ground to diapause, where we could not count them. Data from 2020 were excluded due to inconsistent estimates of the number of surviving webs and larvae entering diapause during the summer of Covid-19.

### Intraguild predation data and ungulate density

We recorded an egg cluster as lost to intraguild predation if the leaf and branch containing the eggs were browsed; intraguild predation was the proportion of egg clusters consumed by ungulates each year. We sourced deer and elk post-hunt population estimates for the area that contained Gothic, Colorado, as reported by Colorado Parks and Wildlife (https://cpw.state.co.us/activities/hunting/big-game). We used ungulate population estimates from the same year that the egg cluster and larval data were collected. We calculated deer or elk density based on the size of the respective hunting unit and the reported population size. We assumed a uniform density of each ungulate across its hunting unit and summed the deer and elk densities to obtain the overall ungulate density metric. We did not examine deer and elk separately because we were solely interested in a total density-dependent effect on intraguild predation.

### Postulated abiotic & biotic drivers

The weather and climate variables we used were based on hypothesized drivers of plant, butterfly, and ungulate traits (Table 1). Both components of pre-diapause survival should be driven by plant water stress and larval growth & metabolic expenditure. Plant water stress should be affected by the previous winter’s accumulation of snow and/or spring and summer precipitation patterns. Larval growth and metabolic expenditure should be altered by daily minimum and maximum temperatures during the growth period. We hypothesized that web survival was driven by intraguild predation. Intraguild predation was expected to be linked to ungulate foraging intensity which should increase with plant resource quality and quantity (e.g. Ruprecht et al. 2020; Merems et al. 2020). Thus, we predicted that spring and summer temperature and precipitation patterns would drive intraguild predation. Intraguild predation should also be driven by human activity in the study site (e.g. Oberosler et al. 2017; Pęksa & Ciach, 2018) and ungulate density.

**Table 1:** Candidate *a priori* predictor set (a) for pre-diapause survival and intraguild predation and (b) retained predictors after filtering for strong Pearson pairwise correlations (|ρ| > 0.5) and variance inflation factors (VIF < 5). Predictors used for model selection were those that passed both filter steps and are bolded. Superscripts “a” and “b” denote predictors retained respectively for web survival or larval survival.

| Response | Hypothesized drivers (a) | Retained predictor set (b) |
| --- | --- | --- |
| Pre-diapause survival | *Mean temperatures:*<br>Daily minimums & maximums: June – August<br>Summer Mean (June, July, August)<br>*Precipitation:*<br>Total summer rainfall (June, July, August)<br>Total snowfall (the previous winter)<br>*Climate indices:*<br>ENSO 3.4, PDO, PNA: Spring (March, April, May) & Summer mean (June, July, August)<br>*Biotic drivers:*<br>Intraguild predation<br>Ungulate population density | *Mean temperatures:*<br>None<br>*Precipitation:*<br>**Summer rainfall<sup>a</sup>**<br>*Climate indices:*<br>**April PNA<sup>b</sup>, Summer PDO<sup>b</sup>, Spring PNA<sup>b</sup>**<br>*Biotic drivers:*<br>**Intraguild Predation<sup>a</sup>** |
| Intraguild Predation | *Mean temperatures:*<br>Daily minimums & maximums: June - August<br>Summer mean (June, July, August)<br>*Precipitation:*<br>Total rainfall: June – August<br>Total precipitation summer (June, July, August)<br>Total snowfall (the previous winter)<br>*Climate indices:*<br>ENSO 3.4, PDO, PNA: Spring (March, April, May) & Summer mean (June, July, August)<br>*Biotic drivers:*<br>Total number of days of researcher activity<br>Ungulate population density | *Mean temperatures:*<br>**June daily minimum**<br>*Precipitation:*<br>**June rainfall**<br>*Climate indices:*<br>**Spring PDO, Summer PDO**<br>*Biotic drivers:*<br>**Researcher activity** |

**Table 2:** Bayesian SEM model estimates (log-odds), estimated error, and credible intervals for factors affecting pre-diapause web & larval survival and intraguild predation. Web survival was fit with the binomial distribution; larval survival and intraguild predation were fit with the beta distribution. Phi is a distributional parameter relevant to the beta distribution. R-hat measures whether each Markov chain produced similar estimates. Bulk effective sample size (ESS) reports the number of draws used to calculate the estimate. Tail ESS reports the number of draws used to calculate the credible interval. Direction probability is a measure of confidence in an estimate.

| **Response**<br>**R<sup>2</sup> [95% CI]** | **Predictor** | **Estimate**<br>**[95% CI]** | **Est.**<br>**Error** | **Bulk**<br>**ESS** | **Tail**<br>**ESS** | **R-hat** | **Direction**<br>**Probability** |
| --- | --- | --- | --- | --- | --- | --- | --- |
| Web<br>Survival<br>0.98 ± 0.002<br>[0.97, 0.98] | Intercept | 1.41<br>[1.30, 1.53] | 0.06 | 86,468 | 52,633 | 1.00 | 100% |
|  | Intraguild<br>Predation | -0.58<br>[-0.79, -0.36] | 0.11 | 80,261 | 51,330 | 1.00 | 100% |
|  | Summer rainfall | -0.34<br>[-0.55, -0.12] | 0.11 | 72,028 | 53,229 | 1.00 | 99% |
| Larval<br>Survival<br>0.56 ± 0.11<br>[0.28, 0.71] | Intercept | -1.08<br>[-1.17, -0.99] | 0.05 | 75,634 | 46,457 | 1.00 | 100% |
|  | Phi | 158.89<br>[75.20, 272.57] | 51.07 | 71,375 | 44,871 | 1.00 |  |
|  | Spring PNA | -0.47<br>[-0.67, -0.27] | 0.10 | 72,610 | 47,404 | 1.00 | 100% |
| Intraguild<br>predation<br>0.68 ± 0.10<br>[0.43, 0.77] | Intercept | -2.56<br>[-2.84, -2.27] | 0.15 | 62,519 | 45,645 | 1.00 | 100% |
|  | Phi | 49.62<br>[22.61, 86.61] | 16.53 | 60,918 | 44,522 | 1.00 | 100% |
|  | June rainfall | 0.96<br>[0.40, 1.49] | 0.28 | 61,402 | 47,068 | 1.00 | 99% |
|  | Spring PDO | -0.83<br>[-1.42, -0.26] | 0.20 | 61,688 | 49,734 | 1.00 | 99% |

We included the El Niño Southern Oscillation (ENSO), Pacific Decadal Oscillation (PDO), and Pacific-North American climate pattern (PNA), because they are associated with precipitation and temperature variability in the western United States (McCabe & Dettinger, 1999; Mantua & Hare, 2002; Leathers et al. 1991). The timing of spring is generally earlier in Colorado during the positive phase of spring PDO and during a spring El Niño; the inverse is true for the negative PDO phase and for a La Niña (McCabe et al. 2012). Likewise, snowmelt is generally earlier in Colorado during the positive phase of spring PNA; the inverse is true for the negative phase (Ballinger et al. 2018).

### Sourced abiotic drivers

Temperature and rainfall data were obtained from the NOAA National Weather Service Station located in Crested Butte, Colorado (www.ncei.noaa.gov), 9.5 km south of the study site. The ENSO 3.4, PDO, and PNA indices were sourced from NOAA’s National Centers for Environmental Information database (www.ncei.noaa.gov). Long-term snow data were collected in Gothic, Colorado (https://www.gothicwx.org), 0.37 km from the study site.

### Statistical analysis

We evaluated the direct and indirect effects of the drivers of pre-diapause survival and intraguild predation via Bayesian structural equation modelling (BSEM). We used a Bayesian SEM because of its robustness in handling small sample sizes (Fan et al. 2016). Prior to fitting models, we filtered the candidate predictor set for each response variable to reduce the parameter space and remove multicollinearity among predictors. We first calculated Pearson correlation coefficients (ρ) between each response variable and its hypothesized predictor set and retained only predictors where |ρ| > 0.5 (Tredennick et al. 2021; Table 1). Ungulate density was weakly correlated (|ρ| < 0.5) with intraguild predation and thus was removed as a predictor. We then obtained the variance inflation factors among the remaining predictors for each response variable to ensure there was no multicollinearity (Dormann et al. 2013; Table 1). Finally, we standardized the remaining predictors by mean centering and dividing by two standard deviations (Gelman, 2008).

We used regularization and Bayesian leave-one-out cross validation (loo-cv) (Hooten & Hobbs, 2016; Yates et al. 2022) to select the most explanatory model for each response variable. Regularization was done using shrinkage priors that we evaluated using pareto-smoothed importance sampling loo-cv (Yates et al. 2022). We used weakly informative priors that effectively shrunk parameter estimates towards zero unless there was strong evidence of a large effect (McElreath, 2015; Lemoine, 2019). We fit multiple models of each response variable with their respective reduced predictor set and a candidate prior. Each model was ranked by expected log predictive density (ELPD) and the loo information criterion (LOOIC). Among the loo-cv results, there was no indication of which prior best described each response variable because they had similar ELPD values (Table S6). Thus, we used posterior predictive checks (Fig. S7 – S9; Gabry et al. 2019) and a prior powerscaling sensitivity analysis (Table S7 – S9; Kallioinen et al. 2025).

We built Bayesian regression models that exhausted all possible additive combinations of the reduced predictor set for each response variable. We ignored interactive combinations to reduce the parameter space. We fit web survival using the binomial family distribution; larval survival and intraguild predation were fit using the beta distribution. We also fit each model with the logit link function, the appropriate shrinkage prior, and used the Bayesian loo-cv resampling method to compare models. We reduced overfitting risk by using the modified one-standard-error rule (m-OSE) of Yates et al. (2022) and the selection-induced bias correction method of McLatchie and Vehtari (2024). The m-OSE reduces estimation uncertainty by favoring the least complex model whose adjusted error includes the estimate from the best-scoring model. We also estimated and corrected for selection induced bias that favored more complex models from our Bayesian loo-cv model comparisons (McLatchie and Vehtari, 2024). We assessed convergence using 4 Markov chains that ran for 10000 iterations after discarding the first 2000 samples. We selected the model that met the m-OSE rule with the highest bias-corrected ELPD and lowest LOOIC. For the larval survival model path, there was no consensus of a top model among the loo-cv results. Thus, we selected the most parsimonious model (Table S2).

Our BSEM was fit using 5 markov chains that ran for 13000 iterations after discarding the first 2000 samples. We assessed BSEM goodness of fit via posterior predictive checks and tests of conditional independence. Conditional independence was used to test if we failed to include any relevant predictor-response relationships in our models (Shipley, 2000). To test each independence claim, we (1) fit separate BSEMs with a claim included, (2) generated model predictions, and (3) compared the overlap coefficient between the predictions from the original model structure and the independence claim.

Statistical analyses were all performed in R version 4.4.3 (R Core Team, 2025). Bayesian regressions and the BSEM were fit using the ‘brms’ R package (Bürkner, 2017). VIFs were calculated using the ‘car’ R package (Fox, 2019). Leave-one-out cross validation was done using the ‘loo’ r package (Vehtari et al. 2024). Bayesian regression diagnostics were done using the ‘bayestestR’ r package (Makowski et al. 2019). Temporal autocorrelation was assessed using the ‘stats’ r package (R Core Team, 2025). Prior and posterior predictive checks were done using the ‘rstanarm’ r package (Goodrich et al. 2024).

## Results

### Direct effects of intraguild predation, weather, and teleconnection patterns on pre-diapause survival

For every increase in intraguild predation, the yearly odds of web survival decreased by 44% (Odds Ratio (OR) = 0.56, 95% CI = [0.45, 0.70]). Increases in the total amount of summer rainfall decreased the odds of web survival by 29% (OR = 0.71, 95% CI = [0.45, 0.70]). Increases in the spring mean of the Pacific-North American pattern decreased the odds of larval survival by 37% (OR = 0.63, 95% CI = [0.58, 0.76]), likely via earlier snowmelt. We performed multiple tests of conditional independence for both web and larval survival. We found that each tested predictor had miniscule effects and credible intervals that overlapped zero (Table S4; Fig. S1 & S2). We found no evidence of a lag-1 autocorrelation in the time series for web survival (acf = -0.26, 95% CI = [-0.54, 0.19]) or larval survival (acf = -0.33, 95% CI = [-0.51, 0.19]).

### Indirect effects of climatic influences on pre-diapause survival via intraguild predation

Weather and a climate teleconnection pattern had indirect effects on pre-diapause survival via intraguild predation. Increases in June rainfall increased the odds of intraguild predation by 161% (OR = 2.61, 95% CI = [1.50, 4.44]), indicating a strong negative indirect effect on web survival (Fig. 2). Increases in the spring mean of the Pacific Decadal Oscillation decreased the odds of intraguild predation by 57% (OR = 0.44, 95% CI = [0.24, 0.77]), likely via earlier spring onset, indicating a positive indirect effect on web survival (Fig. 2). We performed two tests of conditional independence in which we added either summer rainfall or the spring mean of PNA to the best model of intraguild predation. The conditional independence tests revealed that adding either predictor explained little to no effect on intraguild predation and were characterized by credible intervals that overlapped zero (Table S4; Fig. S3). We found no evidence of a lag-1 autocorrelation in the time series of intraguild predation (acf = 0.13, 95% CI = [-0.43, 0.46]).

**Figure 2.**
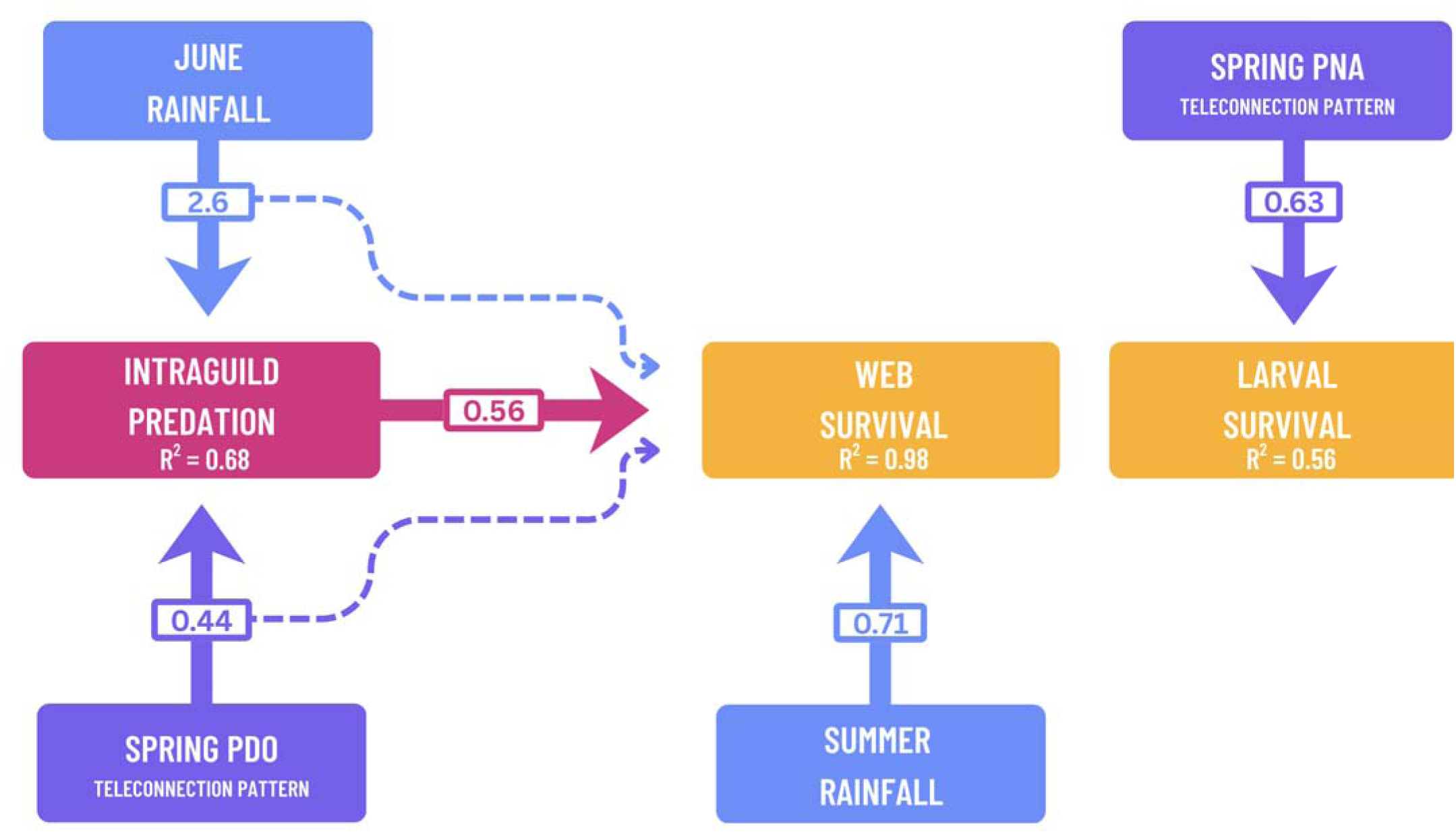
Final Bayesian SEM modeling the relationship between biotic and abiotic factors of pre-diapause survival. PNA is the Pacific North American pattern and PDO is the Pacific Decadal Oscillation. Values contained within boxes represent odds ratios. An odds ratio < 1 describes a negative relationship; An odds ratio > 1 indicates a positive relationship. Arrows represent directional relationships among variables. Direct effects are represented by continuous lines and indirect effects by dashed lines. The thickness of paths has been scaled according to the standardized estimate located within the associated box. Bayesian R^2^’s for the basis set is given in the corresponding boxes.

## Discussion

Theory has long emphasized the ecological significance and ubiquity of intraguild predation across taxa and ecosystems (e.g., Polis et al. 1989; Arim & Marquet, 2004). Despite this, our study is among the first empirical analyses to show the role of ungulate intraguild predation as a driver of insect demography. Additionally, our study addresses an earlier hypothesis that posited teleconnection patterns were better predictors of demographic processes than local weather conditions (Hallet et al. 2004). Here, we found complex effects of both weather and teleconnection patterns on pre-diapause survival. The direction of the effects of teleconnection patterns shifted from negative to positive when mediated through intraguild predation. Finally, our finding that the two components of pre-diapause survival had distinct drivers highlights the importance of careful consideration of the demographic processes behind vital rates. In our system, web survival describes the probability that an egg cluster hatches and the resulting larvae form a hibernaculum at diapause, whereas larval survival describes the survivorship among individual eggs within a cluster. Thus, it is reasonable that both components would be driven by distinct mechanisms despite describing the same life history stage.

### Direct effects of weather and climatic factors on pre-diapause survival

Decreases in pre-diapause web survival were correlated with increases in the total amount of summer rainfall. Such negative effects of increased rainfall throughout the egg and larval development period may be driven by two mechanisms. Rainfall can decrease survivorship simply by increasing egg dislodgement rates (e.g. Rahman et al. 2019; Santos et al. 2020) or, in some cases, by washing off early-instar larvae on to the ground (Bonhof and Overholt, 2001). Eggs and/or early instar larvae that fall to the ground are unlikely to survive given moist, shady, and predator-rich conditions (Williams, 1984). Alternatively, changes in microclimatic conditions following rainfall events have resulted in extended egg and larval development rates leading to decreased probability of survival. For instance, simulated downpours increased larval development rates for two species of Lepidoptera via lower temperatures experienced at leaf surfaces (Chen et al. 2019).

Decreases in pre-diapause larval survival were linked to increases in the spring mean of PNA. Ballinger et al. (2019) showed that decreased spring snowcover extent over the Western United States was associated with positive values of spring PNA. Thus, the effect of spring PNA on larval survival may be driven by a lagged effect of early snowmelt on hostplant quality. Early snowmelt can drive initial patterns of plant water stress via reductions in soil moisture content in montane systems (Harpold, 2016). Additionally, early snowmelt both prolongs and intensifies the period of drought occuring before the onset of the North American Monsoon in our region, thus further driving plant water stress (Sloat et al. 2015). Water stress can alter concentrations of phytohormones and plant secondary metabolites (Lin et al. 2023) which in turn shapes chewing insect herbivore performance, survival, and growth (e.g. Han et al. 2016; Nguyen et al. 2016; Carvajal Acosta et al. 2022). In our region, such effects of plant water stress induced by early snowmelt have been linked to decreases in aphid abundance (Mooney et al. 2021).

### Direct effects of intraguild predation on pre-diapause web survival

We report negative effects of ungulate driven intraguild predation on pre-diapause web survival across an 18-year study. The few studies that have examined this interaction are short-term, do not consider vital rates, and do not attempt to understand its drivers (e.g. Zamora & Gómez, 1993; Gómez & González-Megías, 2002; Bonal & Muñoz, 2007). The impact of intraguild predation is more complex than other biotic interactions (e.g. predation, competition) because it incorporates aspects of predation and exploitative competition (Polis et al. 1989). Thus, our study may understate the degree to which intraguild predation drives insect populations, since we lacked data on resource competition (e.g. hostplant abundance). We used ungulate density as a proxy for competitive effects on pre-diapause web survival but its weak correlation suggests it has little to no effect. Our findings show that even the consumptive aspect of intraguild predation, alone, can be critical to insect population dynamics. Future studies should include data on hostplant abundance to track the competitive aspect of intraguild predation.

### Indirect effects of weather and climatic factors on pre-diapause web survival through intraguild predation

The role of abiotic factors on intraguild predation is generally unclear apart from a single study that found it varied with habitat structure (Finke & Denno, 2002). Here, we found contrasting effects of weather and a teleconnection pattern on intraguild predation. June rainfall promoted intraguild predation likely via a lagged effect of improved plant quality that increased ungulate foraging. Our result is consistent with Ferretti et al.’s (2019) result that increases in rainfall in a mountain meadow had a lagged effect on plant quality that increased alpine ungulate forage intake 45 days later. This suggests that June rainfall could indirectly decrease pre-diapause web survival by increasing intraguild predation in July. Additionally, we found that spring PDO decreased intraguild predation likely via a lagged effect where earlier spring conditions caused ungulates to migrate from winter grounds to the study site sooner. Several studies have shown that ungulate spring migration patterns closely track plant budbreak (e.g. Sawyer & Kauffman, 2011; Merkle et al. 2016; Aikens et al. 2017). This suggests that spring PDO indirectly increased pre-diapause web survival by reducing the overlap between when ungulates forage and when egg clusters develop.

## Conclusion

Our study documents complex effects of local weather and teleconnection patterns on an insect vital rate. The direction of these effects depended on whether they were mediated through intraguild predation. Additionally, we show that seasonal indices of teleconnection patterns can have readily detectable effects on insects, which suggests that seasonal teleconnection effects may be an underexamined demographic driver. Our work expands on an old debate of the relationship between biotic and abiotic drivers – that together they can determine patterns of change in demography. Ultimately, this calls for population ecology to move towards adoption of sequential hypotheses that capture the cascading effects of abiotic factors via biotic interactions.

## Acknowledgements

We thank numerous undergraduate and graduate students for assistance in data collection for this long-term project. Funding came from Stanford University’s Biological Field Studies Program, the University of South Carolina, and the Rocky Mountain Biological Laboratory’s graduate fellowship (to LD) and Navjot Sodhi Conservation Research award (to CLB). We thank members of the Boggs and Ward Watt labs, Alissa Armstrong, Tad Dallas, Eric LoPresti, and Jay Rosenheim for discussion, and Eric LoPresti for comments on the manuscript. The authors recognize the establishment of the mining town of Gothic, which later became the Rocky Mountain Biological Laboratory, as the ancestral homeland of the Uncompahgre (Tabeguache) Ute band. Ute peoples can be considered among the earliest ecologists of the Gunnison Valley.

## References

Aikens, Ellen O., Matthew J. Kauffman, Jerod A. Merkle, Samantha P. H. Dwinnell, Gary L. Fralick, and Kevin L. Monteith. 2017. “The Greenscape Shapes Surfing of Resource Waves in a Large Migratory Herbivore.” Ecology Letters 20 (6): 741–50. 10.1111/ele.12772.

Andrewartha, H. G., and L. C. Birch. 1954. The Distribution and Abundance of Animals. University of Chicago Press.

Arim, Matías, and Pablo A. Marquet. 2004. “Intraguild Predation: A Widespread Interaction Related to Species Biology: Intraguild Predation.” Ecology Letters 7 (7): 557–64. 10.1111/j.1461-0248.2004.00613.x.

Ballinger, T. J., R. V. Rohli, M. J. Allen, D. A. Robinson, and T. W. Estilow. 2018. “Half-Century Perspectives on North American Spring Snowline and Snow Cover Associations with the Pacific-North American Teleconnection Pattern.” Climate Research 74 (3): 201–16. 10.3354/cr01499.

Barr, William B. (2026). Gothic, CO Weather. Available at: https://www.gothicwx.org/. Last accessed 08, February 2026.

Benton, Tim G., Stewart J. Plaistow, and Tim N. Coulson. 2006. “Complex Population Dynamics and Complex Causation: Devils, Details and Demography.” *Proceedings*. Biological Sciences 273 (1591): 1173–81. 10.1098/rspb.2006.3495.

Boggs, Carol L., and Dennis D. Murphy. 1997. “Community Composition in Mountain Ecosystems: Climatic Determinants of Montane Butterfly Distributions.” Global Ecology and Biogeography Letters 6: 39. 10.2307/2997525.

Boggs, Carol L., and David W. Inouye. 2012. “A Single Climate Driver Has Direct and Indirect Effects on Insect Population Dynamics.” Ecology Letters 15 (5): 502–8. 10.1111/j.1461-0248.2012.01766.x.

Bonal, Raúl, and Alberto Muñoz. 2007. “Multi-Trophic Effects of Ungulate Intraguild Predation on Acorn Weevils.” Oecologia 152 (3): 533–40. 10.1007/s00442-007-0672-8.

Bonebrake, Timothy C., Carol L. Boggs, Jessica M. McNally, Jai Ranganathan, and Paul R. Ehrlich. 2010. “Oviposition Behavior and Offspring Performance in Herbivorous Insects: Consequences of Climatic and Habitat Heterogeneity.” Oikos 119 (6): 927–34. 10.1111/j.1600-0706.2009.17759.x.

Bonhof, M. J., and W. A. Overholt. 2001. “Impact of Solar Radiation, Rainfall and Cannibalism on Disappearance of Maize Stemborers in Kenya.” International Journal of Tropical Insect Science 21 (04): 403–7. 10.1017/s1742758400008523.

Bürkner, P. 2017. “Brms: An R Package for Bayesian Multilevel Models Using Stan.” Journal of Statistical Software 080 (August): 1–28. 10.18637/JSS.V080.I01.

Carvajal Acosta, Alma N., Anurag A. Agrawal, and Kailen Mooney. 2023. “Plant Water-use Strategies as Mediators of Herbivore Drought Response: Ecophysiology, Host Plant Quality and Functional Traits.” The Journal of Ecology 111 (3): 687–700. 10.1111/1365-2745.14059

Chen, Cong, Jeffrey A. Harvey, Arjen Biere, and Rieta Gols. 2019. “Rain Downpours Affect Survival and Development of Insect Herbivores: The Specter of Climate Change?” Ecology 100 (11): e02819. 10.1002/ecy.2819.

Coulson, T., E. A. Catchpole, S. D. Albon, B. J. Morgan, J. M. Pemberton, T. H. Clutton-Brock, M. J. Crawley, and B. T. Grenfell. 2001. “Age, Sex, Density, Winter Weather, and Population Crashes in Soay Sheep.” *Science (New York*, N.Y.) 292 (5521): 1528–31. 10.1126/science.292.5521.1528.

Colorado Parks & Wildlife. (2026). Hunting Statistics. Available at: https://cpw.state.co.us/activities/hunting/big-game. Last accessed 08, February 2026.

Dormann, Carsten F., Jane Elith, Sven Bacher, Carsten Buchmann, Gudrun Carl, Gabriel Carré, Jaime R. García Marquéz, et al. 2013. “Collinearity: A Review of Methods to Deal with It and a Simulation Study Evaluating Their Performance.” Ecography 36 (1): 27–46. 10.1111/j.1600-0587.2012.07348.x.

Duncan, Rebecca J., Margaret E. Andrew, and Mads C. Forchhammer. 2021. “Snow Mediates Climatic Impacts on Arctic Herbivore Populations.” Polar Biology 44 (7): 1251–71. 10.1007/s00300-021-02871-y.

Elton, Charles. 1949. “Population Interspersion: An Essay on Animal Community Patterns.” The Journal of Ecology 37 (1): 1. 10.2307/2256726.

Fan, Yi, Jiquan Chen, Gabriela Shirkey, Ranjeet John, Susie R. Wu, Hogeun Park, and Changliang Shao. 2016. “Applications of Structural Equation Modeling (SEM) in Ecological Studies: An Updated Review.” Ecological Processes 5 (1). 10.1186/s13717-016-0063-3.

Ferretti, F., S. Lovari, and P. Stephens. 2018. “Joint Effects of Weather and Interspecific Competition on Foraging Behavior and Survival of a Mountain Herbivore.” Current Zoology 65 (May): 165–75. 10.1093/cz/zoy032.

Finke, D. L., and R. F. Denno. 2003. “Intra-Guild Predation Relaxes Natural Enemy Impacts on Herbivore Populations.” Ecological Entomology 28 (1): 67–73. 10.1046/j.1365-2311.2003.00475.x.

Finke, Deborah L., and Robert F. Denno. 2002. “Intraguild Predation Diminished in Complex-Structured Vegetation: Implications for Prey Suppression.” Ecology 83 (3): 643. 10.2307/3071870.

Fox, John, Sanford Weisberg, and Brad Price. 2019. An R Companion to Applied Regression. 3rd ed. Sage. https://www.john-fox.ca/Companion/.

Frederiksen, Morten, Sarah Wanless, Michael P. Harris, Peter Rothery, and Linda J. Wilson. 2004. “The Role of Industrial Fisheries and Oceanographic Change in the Decline of North Sea Black-legged Kittiwakes: Kittiwake Decline: Fishery or Oceanography?” The Journal of Applied Ecology 41 (6): 1129–39. 10.1111/j.0021-8901.2004.00966.x.

Gabry, Jonah, Daniel Simpson, Aki Vehtari, Michael Betancourt, and Andrew Gelman. 2019. “Visualization in Bayesian Workflow.” *Journal of the Royal Statistical Society. Series A*, (Statistics in Society) 182 (2): 389–402. 10.1111/rssa.12378.

Gelman, Andrew. 2008. “Scaling Regression Inputs by Dividing by Two Standard Deviations.” Statistics in Medicine 27 (15): 2865–73. 10.1002/sim.3107.

Gish, Moshe, Matan Ben-Ari, and Moshe Inbar. 2017. “Direct Consumptive Interactions between Mammalian Herbivores and Plant-Dwelling Invertebrates: Prevalence, Significance, and Prospectus.” Oecologia 183 (2): 347–52. 10.1007/s00442-016-3775-2.

Gómez, José M., and Adela González-Megías. 2002. “Asymmetrical Interactions between Ungulates and Phytophagous Insects: Being Different Matters.” Ecology 83 (1): 203–11. 10.1890/0012-9658(2002)083%5B0203:aibuap%5D2.0.co;2.

Goodrich Ben, Jonah Gabry, Imad Ali, and Sam Brilleman. (2025). Rstanarm: Bayesian applied regression modeling via Stan. R package version 2.32.1. https://mc-stan.org/rstanarm.

Goodsman, Devin W., Guenchik Grosklos, Brian H. Aukema, Caroline Whitehouse, Katherine P. Bleiker, Nate G. McDowell, Richard S. Middleton, and Chonggang Xu. 2018. “The Effect of Warmer Winters on the Demography of an Outbreak Insect Is Hidden by Intraspecific Competition.” Global Change Biology 24 (8): 3620–28. 10.1111/gcb.14284.

Govindarajulu, Purnima P., and Bradley R. Anholt. 2006. “Interaction between Biotic and Abiotic Factors Determines Tadpole Survival Rate under Natural Conditions.” Ecoscience 13 (3): 413–21. 10.2980/i1195-6860-13-3-413.1.

Hallett, T. B., T. Coulson, J. G. Pilkington, T. H. Clutton-Brock, J. M. Pemberton, and B. T. Grenfell. 2004. “Why Large-Scale Climate Indices Seem to Predict Ecological Processes Better than Local Weather.” Nature 430 (6995): 71–75. 10.1038/nature02708.

Han, Peng, Nicolas Desneux, Thomas Michel, Jacques Le Bot, Aurelie Seassau, Eric Wajnberg, Edwige Amiens-Desneux, and Anne-Violette Lavoir. 2016. “Does Plant Cultivar Difference Modify the Bottom-up Effects of Resource Limitation on Plant-Insect Herbivore Interactions?” Journal of Chemical Ecology 42 (12): 1293–1303. 10.1007/s10886-016-0795-7.

Harpold, Adrian A. 2016. “Diverging Sensitivity of Soil Water Stress to Changing Snowmelt Timing in the Western U.S.” Advances in Water Resources 92 (June): 116–29. 10.1016/j.advwatres.2016.03.017.

Hindle, Bethan J., Jill G. Pilkington, Josephine M. Pemberton, and Dylan Z. Childs. 2019. “Cumulative Weather Effects Can Impact across the Whole Life Cycle.” Global Change Biology 25 (10): 3282–93. 10.1111/gcb.14742.

Holdren, Cheryl E., and Paul R. Ehrlich. 1981. “Long Range Dispersal in Checkerspot Butterflies: Transplant Experiments with Euphydryas Gillettii.” Oecologia 50 (1): 125–29. 10.1007/BF00378805.

Holt, Robert D., and Gary A. Polis. 1997. “A Theoretical Framework for Intraguild Predation.” The American Naturalist 149 (4): 745–64. 10.1086/286018.

Hone, Jim, and Tim H. Clutton-Brock. 2007. “Climate, Food, Density and Wildlife Population Growth Rate.” The Journal of Animal Ecology 76 (2): 361–67. 10.1111/j.1365-2656.2006.01200.x.

Hooten, M. B., and N. T. Hobbs. 2015. “A Guide to Bayesian Model Selection for Ecologists.” Ecological Monographs 85 (1): 3–28. 10.1890/14-0661.1.

Hunter, M. D., G. C. Varley, and G. R. Gradwell. 1997. “Estimating the Relative Roles of Top-down and Bottom-up Forces on Insect Herbivore Populations: A Classic Study Revisited.” Proceedings of the National Academy of Sciences of the United States of America 94 (17): 9176–81. 10.1073/pnas.94.17.9176.

Juhasz, Claire-Cécile, Bill Shipley, Gilles Gauthier, Dominique Berteaux, and Nicolas Lecomte. 2020. “Direct and Indirect Effects of Regional and Local Climatic Factors on Trophic Interactions in the Arctic Tundra.” The Journal of Animal Ecology 89 (3): 704–15. 10.1111/1365-2656.13104.

Kallioinen, Noa, Topi Paananen, Paul-Christian Bürkner, and Aki Vehtari. 2024. “Detecting and Diagnosing Prior and Likelihood Sensitivity with Power-Scaling.” Statistics and Computing 34 (1). 10.1007/s11222-023-10366-5.

Kingsolver, Joel G. 1989. “Weather and the Population Dynamics of Insects: Integrating Physiological and Population Ecology.” Physiological Zoology 62 (2): 314–34. 10.1086/physzool.62.2.30156173.

Laczi, Miklós, László Zsolt Garamszegi, Gergely Hegyi, Márton Herényi, Gábor Ilyés, Réka Könczey, Gergely Nagy, et al. 2019. “Teleconnections and Local Weather Orchestrate the Reproduction of Tit Species in the Carpathian Basin.” Journal of Avian Biology 50 (12). 10.1111/jav.02179.

Leathers, D., Brent Yarnal, and M. Palecki. 1991. “The Pacific/North American Teleconnection Pattern and United States Climate. Part I: Regional Temperature and Precipitation Associations.” Journal of Climate 4 (May): 517–28. 10.1175/1520-0442(1991)004%3C0517:TPATPA%3E2.0.CO;2.

Lemoine, Nathan P. 2019. “Moving beyond Noninformative Priors: Why and How to Choose Weakly Informative Priors in Bayesian Analyses.” *Oikos (Copenhagen*, Denmark) 128 (7): 912–28. 10.1111/oik.05985.

Lin, Po-An, Jessica Kansman, Wen-Po Chuang, Christelle Robert, Matthias Erb, and Gary W. Felton. 2023. “Water Availability and Plant-Herbivore Interactions.” Journal of Experimental Botany 74 (9): 2811–28. 10.1093/jxb/erac481.

LoPresti, Eric, Billy Krimmel, and Ian S. Pearse. 2018. “Entrapped Carrion Increases Indirect Plant Resistance and Intra-guild Predation on a Sticky Tarweed.” *Oikos (Copenhagen*, Denmark) 127 (7): 1033–44. 10.1111/oik.04806.

Louthan, Allison M., Jeffrey R. Walters, Adam J. Terando, Victoria Garcia, and William F. Morris. 2021. “Shifting Correlations among Multiple Aspects of Weather Complicate Predicting Future Demography of a Threatened Species.” *Ecosphere (Washington*, D.C) 12 (9). 10.1002/ecs2.3740.

Mabille, Géraldine, Sébastien Descamps, and Dominique Berteaux. 2010. “Predation as a Probable Mechanism Relating Winter Weather to Population Dynamics in a North American Porcupine Population.” Population Ecology 52 (4): 537–46. 10.1007/s10144-010-0198-5.

Makowski, Dominique, Mattan Ben-Shachar, and Daniel Lüdecke. 2019. “BayestestR: Describing Effects and Their Uncertainty, Existence and Significance within the Bayesian Framework.” Journal of Open Source Software 4 (40): 1541. 10.21105/joss.01541.

Mantua, Nathan J., and Steven R. Hare. 2002. “The Pacific Decadal Oscillation.” Journal of Oceanography 58 (1): 35–44. 10.1023/a:1015820616384.

Marini, Lorenzo, Åke Lindelöw, Anna Maria Jönsson, Sören Wulff, and Leif Martin Schroeder. 2013. “Population Dynamics of the Spruce Bark Beetle: A Long-term Study.” *Oikos (Copenhagen*, Denmark) 122 (12): 1768–76. 10.1111/j.1600-0706.2013.00431.x.

Marshal, Jason P., Paul R. Krausman, and Vernon C. Bleich. 2005. “Rainfall, Temperature, and Forage Dynamics Affect Nutritional Quality of Desert Mule Deer Forage.” Rangeland Ecology & Management 58 (4): 360–65. 10.2111/1551-5028(2005)058%5B0360:rtafda%5D2.0.co;2.

McCabe, G., T. Ault, B. Cook, J. Betancourt, and M. D. Schwartz. 2012. “Influences of the El Niño Southern Oscillation and the Pacific Decadal Oscillation on the Timing of the North American Spring.” International Journal of Climatology 32 (GSFC-E-DAA-TN8995). 10.1002/joc.3400.

McCabe, Gregory J., and Michael D. Dettinger. 1999. “Decadal Variations in the Strength of ENSO Teleconnections with Precipitation in the Western United States.” International Journal of Climatology: A Journal of the Royal Meteorological Society 19 (13): 1399–1410. 10.1002/(sici)1097-0088(19991115)19:13%3C1399::aid-joc457%3E3.0.co;2-a.

McElreath, Richard. 2015. Statistical Rethinking: A Bayesian Course with Examples in R and Stan. London, England: Chapman & Hall/CRC.

McLatchie, Yann, and Aki Vehtari. 2024. “Efficient Estimation and Correction of Selection-Induced Bias with Order Statistics.” Statistics and Computing 34 (4): 132. 10.1007/s11222-024-10442-4.

Merems, Jennifer L., Lisa A. Shipley, Taal Levi, Joel Ruprecht, Darren A. Clark, Michael J. Wisdom, Nathan J. Jackson, Kelley M. Stewart, and Ryan A. Long. 2020. “Nutritional-Landscape Models Link Habitat Use to Condition of Mule Deer (Odocoileus Hemionus).” Frontiers in Ecology and Evolution 8 (April). 10.3389/fevo.2020.00098.

Merkle, Jerod A., Kevin L. Monteith, Ellen O. Aikens, Matthew M. Hayes, Kent R. Hersey, Arthur D. Middleton, Brendan A. Oates, Hall Sawyer, Brandon M. Scurlock, and Matthew J. Kauffman. 2016. “Large Herbivores Surf Waves of Green-up during Spring.” *Proceedings*. Biological Sciences 283 (1833): 20160456. 10.1098/rspb.2016.0456.

Mooney, Emily, Maria Mullins, James Den Uyl, Samantha Trail, Phuong Nguyen, Janel Owens, Elsa Godtfredsen, and Shane Heschel. 2021. “Early Snowmelt Reduces Aphid Abundance (Aphis Asclepiadis) by Creating Water-Stressed Host Plants (Ligusticum Porteri) and Altering Interactions with Ants.” Arthropod-Plant Interactions 15 (1): 33–46. 10.1007/s11829-020-09793-2.

Morris, William F., Catherine A. Pfister, Shripad Tuljapurkar, Chirrakal V. Haridas, Carol L. Boggs, Mark S. Boyce, Emilio M. Bruna, et al. 2008. “Longevity Can Buffer Plant and Animal Populations against Changing Climatic Variability.” Ecology 89 (1): 19–25. 10.1890/07-0774.1.

National Oceanic and Atmospheric Administration. (2026). NCDC Climate Data Online NOAA National Centers for Environmental Information. Available at: http://www.ncdc.noaa.gov/. Last accessed 08, February 2026.

Nguyen, Duy, Nunzio D’Agostino, Tom O. G. Tytgat, Pulu Sun, Tobias Lortzing, Eric J. W. Visser, Simona M. Cristescu, et al. 2016. “Drought and Flooding Have Distinct Effects on Herbivore-Induced Responses and Resistance in Solanum Dulcamara: ‘Effects of Drought and Flooding on Insect Defence.’” Plant, Cell & Environment 39 (7): 1485–99. 10.1111/pce.12708.

Nicholson, A. J. 1954. “An Outline of the Dynamics of Animal Populations.” Australian Journal of Zoology 2 (1): 9–65. 10.1071/zo9540009.

Oberosler, Valentina, Claudio Groff, Aaron Iemma, Paolo Pedrini, and Francesco Rovero. 2017. “The Influence of Human Disturbance on Occupancy and Activity Patterns of Mammals in the Italian Alps from Systematic Camera Trapping.” Zeitschrift Für Saugetierkunde [Mammalian Biology] 87 (November): 50–61. 10.1016/j.mambio.2017.05.005.

Ottersen, Geir, Benjamin Planque, Andrea Belgrano, Eric Post, Philip C. Reid, and Nils C. Stenseth. 2001. “Ecological Effects of the North Atlantic Oscillation.” Oecologia 128 (1): 1–14. 10.1007/s004420100655.

Pęksa, Łukasz, and Michał Ciach. 2018. “Daytime Activity Budget of an Alpine Ungulate (Tatra Chamois Rupicapra Rupicapra Tatrica): Influence of Herd Size, Sex, Weather and Human Disturbance.” Mammal Research 63 (4): 443–53. 10.1007/s13364-018-0376-y.

Pol, Martijn van de, Liam D. Bailey, Nina McLean, Laurie Rijsdijk, Callum R. Lawson, and Lyanne Brouwer. 2016. “Identifying the Best Climatic Predictors in Ecology and Evolution.” Methods in Ecology and Evolution 7 (10): 1246–57. 10.1111/2041-210x.12590.

Polis, G. A., C. A. Myers, and R. D. Holt. 1989. “The Ecology and Evolution of Intraguild Predation: Potential Competitors That Eat Each Other.” Annual Review of Ecology and Systematics 20 (1): 297–330. 10.1146/annurev.es.20.110189.001501.

Price, P., C. E. Bouton, P. Gross, B. McPheron, J. Thompson, and A. E. Weis. 1980. “Interactions among Three Trophic Levels: Influence of Plants on Interactions between Insect Herbivores and Natural Enemies.” Annual Review of Ecology, Evolution, and Systematics 11 (November): 41–65. 10.1146/ANNUREV.ES.11.110180.000353.

R Core Team. (2025). R: A Language and Environment for Statistical Computing. Vienna, Austria: R Foundation for Statistical Computing. https://www.R-project.org/.

Rahman, Md Mahbubur, Myron P. Zalucki, and Michael J. Furlong. 2019. “Diamondback Moth Egg Susceptibility to Rainfall: Effects of Host Plant and Oviposition Behavior.” Entomologia Experimentalis et Applicata 167 (8): 701–12. 10.1111/eea.12816.

Roland, Jens, and Stephen F. Matter. 2013. “Variability in Winter Climate and Winter Extremes Reduces Population Growth of an Alpine Butterfly.” Ecology 94 (1): 190–99. 10.1890/12-0611.1.

Rosenheim, Jay A., Lawrence R. Wilhoit, and Christine A. Armer. 1993. “Influence of Intraguild Predation among Generalist Insect Predators on the Suppression of an Herbivore Population.” Oecologia 96 (3): 439–49. 10.1007/BF00317517.

Ruprecht, Joel S., David N. Koons, Kent R. Hersey, N. Thompson Hobbs, and Daniel R. MacNulty. 2020. “The Effect of Climate on Population Growth in a Cold-adapted Ungulate at Its Equatorial Range Limit.” *Ecosphere (Washington*, D.C) 11 (2). 10.1002/ecs2.3058.

Sandvik, H., K. E. Erikstad, and B. E. Sæther. 2012. “Climate Affects Seabird Population Dynamics Both via Reproduction and Adult Survival.” Marine Ecology Progress Series 454: 273–84. https://www.int-res.com/abstracts/meps/v454/p273-284.

Sandvik, Hanno, and Kjell Einar Erikstad. 2008. “Seabird Life Histories and Climatic Fluctuations: A Phylogenetic-comparative Time Series Analysis of North Atlantic Seabirds.” Ecography 31 (1): 73–83. 10.1111/j.2007.0906-7590.05090.x.

Santos, Abraão A., Arthur V. Ribeiro, Scott V. C. Groom, Elizeu S. Farias, Daiane G. Carmo, Renata C. Santos, and Marcelo C. Picanço. 2020. “Season and Weather Affect the Mortality of Immature Stages of*Ascia MonusteOrseis*(Lepidoptera: Pieridae) Caused by Natural Factors.” Austral Entomology 59 (4): 810–18. 10.1111/aen.12500.

Sawyer, Hall, and Matthew J. Kauffman. 2011. “Stopover Ecology of a Migratory Ungulate: *Ungulate Stopover Ecology*.” The Journal of Animal Ecology 80 (5): 1078–87. 10.1111/j.1365-2656.2011.01845.x.

Shipley, Bill. 2000. “A New Inferential Test for Path Models Based on Directed Acyclic Graphs.” Structural Equation Modeling: A Multidisciplinary Journal 7 (2): 206–18. 10.1207/s15328007sem0702_4.

Sloat, Lindsey L., Amanda N. Henderson, Christine Lamanna, and Brian J. Enquist. 2015. “The Effect of the Foresummer Drought on Carbon Exchange in Subalpine Meadows.” *Ecosystems (New York*, N.Y.) 18 (3): 533–45. 10.1007/s10021-015-9845-1.

Stenseth, Nils Chr, Geir Ottersen, James W. Hurrell, Atle Mysterud, Mauricio Lima, Kung-Sik Chan, Nigel G. Yoccoz, and Bjørn Adlandsvik. 2003. “Studying Climate Effects on Ecology through the Use of Climate Indices: The North Atlantic Oscillation, El Niño Southern Oscillation and Beyond.” *Proceedings*. Biological Sciences 270 (1529): 2087–96. 10.1098/rspb.2003.2415.

Tredennick, Andrew T., Giles Hooker, Stephen P. Ellner, and Peter B. Adler. 2021. “A Practical Guide to Selecting Models for Exploration, Inference, and Prediction in Ecology.” Ecology 102 (6): e03336. 10.1002/ecy.3336.

Vehtari, Aki, Andrew Gelman, Jonah Gabry, and Yuling Yao. 2021. “Package ‘Loo.’” Efficient Leave-One-out Cross-Validation and WAIC for Bayesian Models. https://ftp.sun.ac.za/ftp/CRAN/web/packages/loo/refman/loo.html.

Wan, Xinru, Marcel Holyoak, Chuan Yan, Yvon Le Maho, Rodolfo Dirzo, Charles J. Krebs, Nils Chr Stenseth, and Zhibin Zhang. 2022. “Broad-Scale Climate Variation Drives the Dynamics of Animal Populations: A Global Multi-Taxa Analysis.” Biological Reviews of the Cambridge Philosophical Society 97 (6): 2174–94. 10.1111/brv.12888.

White, T. C. R. 2008. “The Role of Food, Weather and Climate in Limiting the Abundance of Animals.” Biological Reviews of the Cambridge Philosophical Society 83 (3): 227–48. 10.1111/j.1469-185X.2008.00041.x.

Williams, Ernest H., Cheryl E. Holdren, and Paul R. Ehrlich. 1984. “The Life History and Ecology of Euphydryas Gillettii Barnes (Nymphalidae).” *Journal of the Lepidopterists’* Society 38 (1): 1–12. https://images.peabody.yale.edu/lepsoc/jls/1980s/1984/1984-38(1)1-Williams.pdf.

Yang, Louie H. 2020. “Toward a More Temporally Explicit Framework for Community Ecology.” Ecological Research 35 (3): 445–62. 10.1111/1440-1703.12099.

Yates, Luke A., Zach Aandahl, Shane A. Richards, and Barry W. Brook. 2023. “Cross Validation for Model Selection: A Review with Examples from Ecology.” Ecological Monographs 93 (1). 10.1002/ecm.1557.

Zamora, R., J. M. Gómez, and J. M. Gómez. 1993. “Vertebrate Herbivores as Predators of Insect Herbivores: An Asymmetrical Interaction Mediated by Size Differences.” *Oikos (Copenhagen*, Denmark) 66 (2): 223. 10.2307/3544808.

