## Supplemental tables 1-9 and Supplemental Figures 1 - 11 for "Intraguild predation, weather, and climate teleconnection patterns interact to determine an insect vital rate"

**Appendix A: Secondary Results**

**Table S1.** All seven models of web survival ranked by expected log predictive density (ELPD), the effective number of parameters (P_loo), and the leave-one-out information criterion (LOOIC). Corrected ELPD was calculated using the selection-induced bias correction from McLatchie and Vehtari (2024). Within OSE was calculated using the modified one-standard-error (m-OSE) rule from Yates et al. (2022). The model selected by the m-OSE is the least complex model whose adjusted error interval includes the estimate of the best-scoring model.

| **Formula** | **ELPD** | **ELPD** | **Δ ELPD** | **P_loo** | **looic** | **Within** |
| --- | --- | --- | --- | --- | --- | --- |
|  | **Corrected** | **(SE)** | **(SE)** | **(SE)** | **(SE)** | **OSE** |
| Intraguild Predation + Summer Rainfall | -66.2 | -66.2 (9.8) |  | 7.5 (2.9) | 132.4  (19.7) | True |
| Intraguild Predation | -77.7 | -69.5 (11.5) | -3.3  (3.5) | 6.4 (2.6) | 139.1 (23.0) | False |
| Intraguild Predation + Summer Rainfall + Apri Snowfall | -78.0 | -69.8 (12.3) | -3.6 (3.2) | 12.2 (4.8) | 139.7  (24.5) | False |
| Intraguild Predation + April Snowfall | -80.4 | -72.2 (15.0) | -6.0 (6.6) | 11.3 (4.9) | 144.4  (30.0) | False |
| Summer Rainfall + April Snowfall | -80.8 | -80.8 (18.7) | -14.6 (10.7) | 13.6 (6.0) | 161.6  (37.4) | False |
| Summer Rainfall | -81.2 | -81.2 (14.2) | -15.0 (8.3) | 10.0 (4.0) | 162.3  (28.3) | False |
| April Snowfall | -83.4 | -83.4 (23.2) | -17.2 (14.5) | 10.4 (5.6) | 166.7 (46.4) | False |

**Table S2.** All seven models of larval survival ranked by expected log predictive density (ELPD), the effective number of parameters (P_loo), and the leave-one-out information criterion (LOOIC). Corrected ELPD was calculated using the selection-induced bias correction from McLatchie and Vehtari (2024). Within OSE was calculated using the modified one-standard-error (m-OSE) rule from Yates et al. (2022). The model selected by the m-OSE is the least complex model whose adjusted error interval includes the estimate of the best-scoring model.

| **Formula** | **ELPD** | **ELPD** | **Δ ELPD** | **P_loo** | **looic** | **Within** |
| --- | --- | --- | --- | --- | --- | --- |
|  | **Corrected** | **(SE)** | **(SE)** | **(SE)** | **(SE)** | **OSE** |
| Spring PNA | 30.1 | 30.1 (2.5) |  | 2.5 (0.7) | -60.2  (5.0) | True |
| Spring PNA + Summer PDO | 27.7 | 29.7 (3.5) | -0.4  (2.0) | 3.9 (1.7) | -59.4 (7.1) | True |
| Spring PNA + April PNA | 27.2 | 29.2 (2.4) | -0.9  (0.7) | 3.5 (0.8) | -58.4  (4.8) | False |
| Spring PNA + April PNA + Summer PDO | 26.7 | 28.7 (3.2) | -1.4 (1.7) | 4.6 (1.5) | -57.5  (6.4) | False |
| April PNA | 26.1 | 26.1 (3.5) | -4.0 (2.8) | 2.7 (0.9) | -52.2  (7.0) | False |
| April PNA + Summer PDO | 25.7 | 25.7 (4.5) | -4.4 (3.6) | 4.1 (1.7) | -51.5  (9.0) | False |
| Summer PDO | 24.5 | 24.7 (5.2) | -5.6 (4.2) | 3.5 (1.8) | -49.0 (10.3) | False |

**Table S3.** Top ten models of Intraguild Predation ranked by expected log predictive density (ELPD), the effective number of parameters (P_loo), and the leave-one-out information criterion (LOOIC). Corrected ELPD was calculated using the selection-induced bias correction from McLatchie and Vehtari (2024). Within OSE was calculated using the modified one-standard-error (m-OSE) rule from Yates et al. (2022). The model selected by the m-OSE is the least complex model whose adjusted error interval includes the estimate of the best-scoring model.

| **Formula** | **ELPD** | **ELPD** | **Δ ELPD** | **P_loo** | **looic** | **Within** |
| --- | --- | --- | --- | --- | --- | --- |
|  | **Corrected** | **(SE)** | **(SE)** | **(SE)** | **(SE)** | **OSE** |
| June Rainfall + Spring PDO | 29.4 | 29.4 (6.3) |  | 4.4 (2.3) | -58.7  (12.7) | True |
| June Rainfall + Spring PDO + June Mean Daily Min Temps | 25.9 | 28.7 (6.2) | -0.7  (0.7) | 4.9 (2.1) | -57.4 (12.3) | False |
| Researcher Activity + June Rainfall + Spring PDO | 25.7 | 28.6 (6.2) | -0.8  (0.5) | 4.9 (2.2) | -57.1  (12.3) | False |
| Spring PDO + June Mean Daily Min Temps | 25.7 | 28.6 (6.0) | -0.8 (1.4) | 4.7 (2.0) | -57.2  (12.0) | False |
| Researcher Activity + Spring PDO + June Mean Daily Min Temps | 25.1 | 27.9 (5.9) | -1.4 (1.3) | 5.2 (2.0) | -55.8  (11.9) | False |
| Researcher Activity + June Rainfall + Spring PDO + June mean daily min temps | 25.0 | 27.8 (6.1) | -1.5 (0.8) | 5.6 (2.2) | -55.6  (12.2) | False |
| Researcher Activity + June Rainfall | 24.8 | 27.6 (4.4) | -1.7 (3.0) | 3.7 (1.1) | -55.3 (8.7) | False |
| June Rainfall | 24.6 | 27.5 (3.8) | -1.9 (4.0) | 2.6 (0.7) | -54.9  (7.4) | False |
| Researcher Activity + June Rainfall + June Mean Daily Min Temps | 23.7 | 26.5 (4.2) | -2.8 (3.3) | 4.6 (1.1) | -53.0  (8.5) | False |
| June Rainfall + June Mean Daily Min Temps | 23.6 | 26.4 (3.7) | -2.9 (4.0) | 3.7 (0.8) | -52.8 (7.4) | False |

**Table S4.** Tests of conditional independence (**in bold**) for web survival, larval survival, and intraguild predation. Estimates represent odds ratios. Bulk effective sample size (ESS) is a measure of the number of draws used to calculate the estimate. Tail ESS is a measure of the number of draws used to calculate the credible interval. The overlap coefficient measures the similarity between the predictions from the original SEM of each response variable and each alternative SEM with a particular independence claim. Direction probability is a measure of confidence in an estimate.

| **Independence claim** | **Estimate** | **Est.** | **95% CI** | | **Bulk** | **Tail** | **Overlap** | **Direction** |
| --- | --- | --- | --- | --- | --- | --- | --- | --- |
|  |  | **Error** | **LL** | **UL** | **ESS** | **ESS** | **Coefficient** | **Probability** |
| Web survival ~ |  |  |  |  |  |  |  |  |
| Intraguild predation + Summer rainfall + **spring PNA** | -0.01 | 0.13 | -0.26 | 0.25 | 73,742 | 53,454 | 96% | 52% |
| Intraguild predation + Summer rainfall + **June rainfall** | 0.09 | 0.22 | -0.33 | 0.53 | 47,320 | 46,528 | 97% | 66% |
| Intraguild predation + Summer rainfall + **Spring PDO** | -0.09 | 0.17 | -0.41 | 0.24 | 45,281 | 50,583 | 95% | 70% |
| Larval survival ~ |  |  |  |  |  |  |  |  |
| Spring PNA + **Intraguild Predation** | -0.07 | 0.11 | -0.29 | 0.16 | 58,195 | 46,400 | 94% | 73% |
| Spring PNA + **Summer rainfall** | 0.00 | 0.10 | -0.19 | 0.20 | 78,916 | 47,988 | 95% | 52% |
| Spring PNA + **June rainfall** | -0.05 | 0.10 | -0.26 | 0.15 | 68,082 | 48,489 | 96% | 70% |
| Spring PNA + **Spring PDO** | -0.04 | 0.12 | -0.28 | 0.20 | 55,173 | 47,291 | 95% | 62% |
| Intraguild Predation ~ |  |  |  |  |  |  |  |  |
| June rainfall + spring PDO + **Summer rainfall** | 0.03 | 0.33 | -0.65 | 0.66 | 45,023 | 41,666 | 96% | 55% |
| June rainfall + spring PDO + **Spring PNA** | -0.09 | 0.34 | -0.78 | 0.56 | 55,035 | 45,907 | 95% | 60% |

**Figure S1.** Density overlap plots of model predictions generated from the Bayesian structural equation model (SEM) for web survival and its respective conditional independence tests. The SEM was defined by the following model structure shown in blue: Web survival ~ IGP + Summer Rainfall. Yellow shows the addition of the tested independence claim to the model.

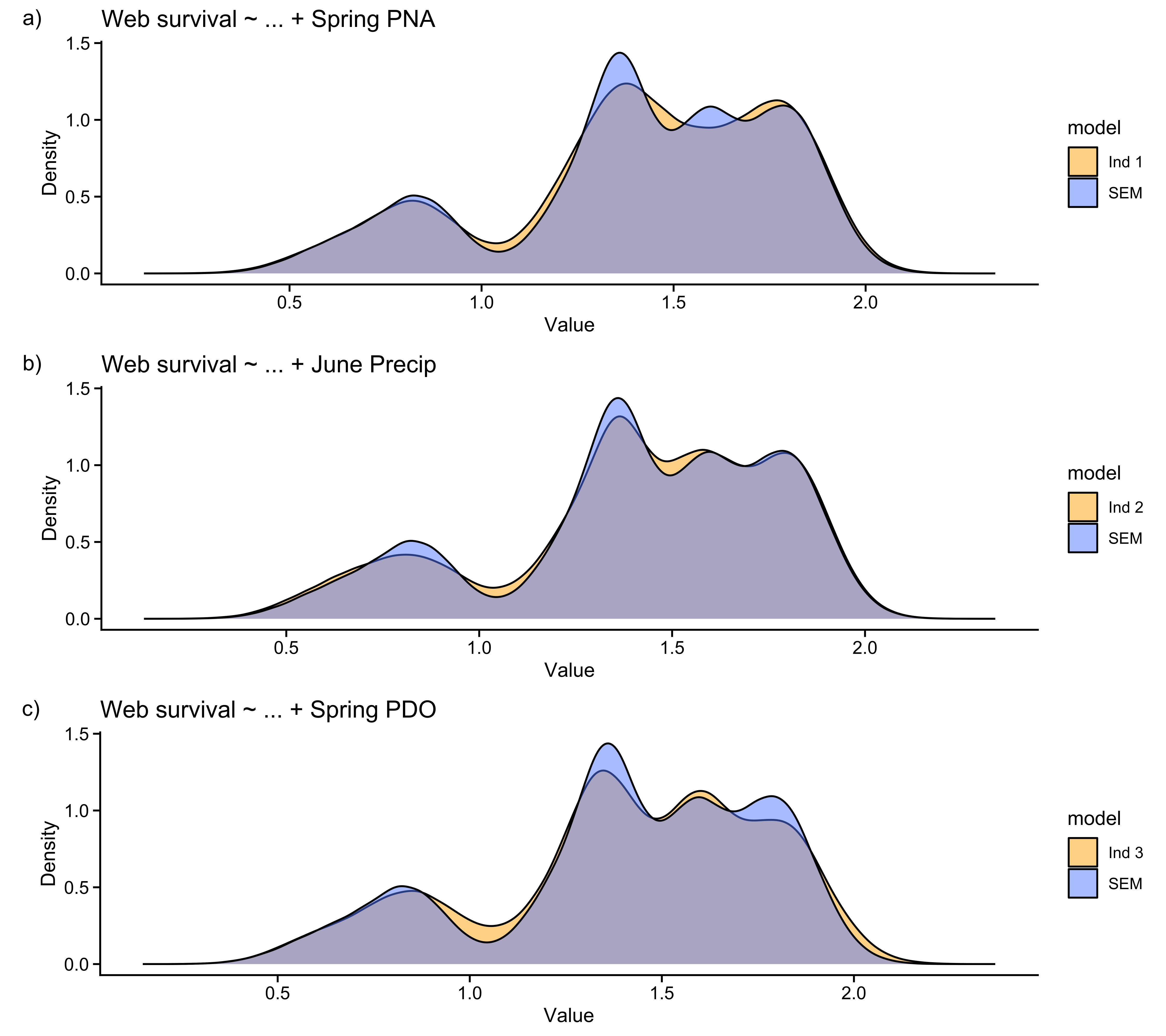

**Figure S2.** Density overlap plots of model predictions generated from the Bayesian structural equation model (SEM) for larval survival and its respective conditional independence tests. The SEM was defined by the following model structure shown in blue: Larval survival ~ spring PNA. Yellow shows the addition of the tested independence claim to the model.

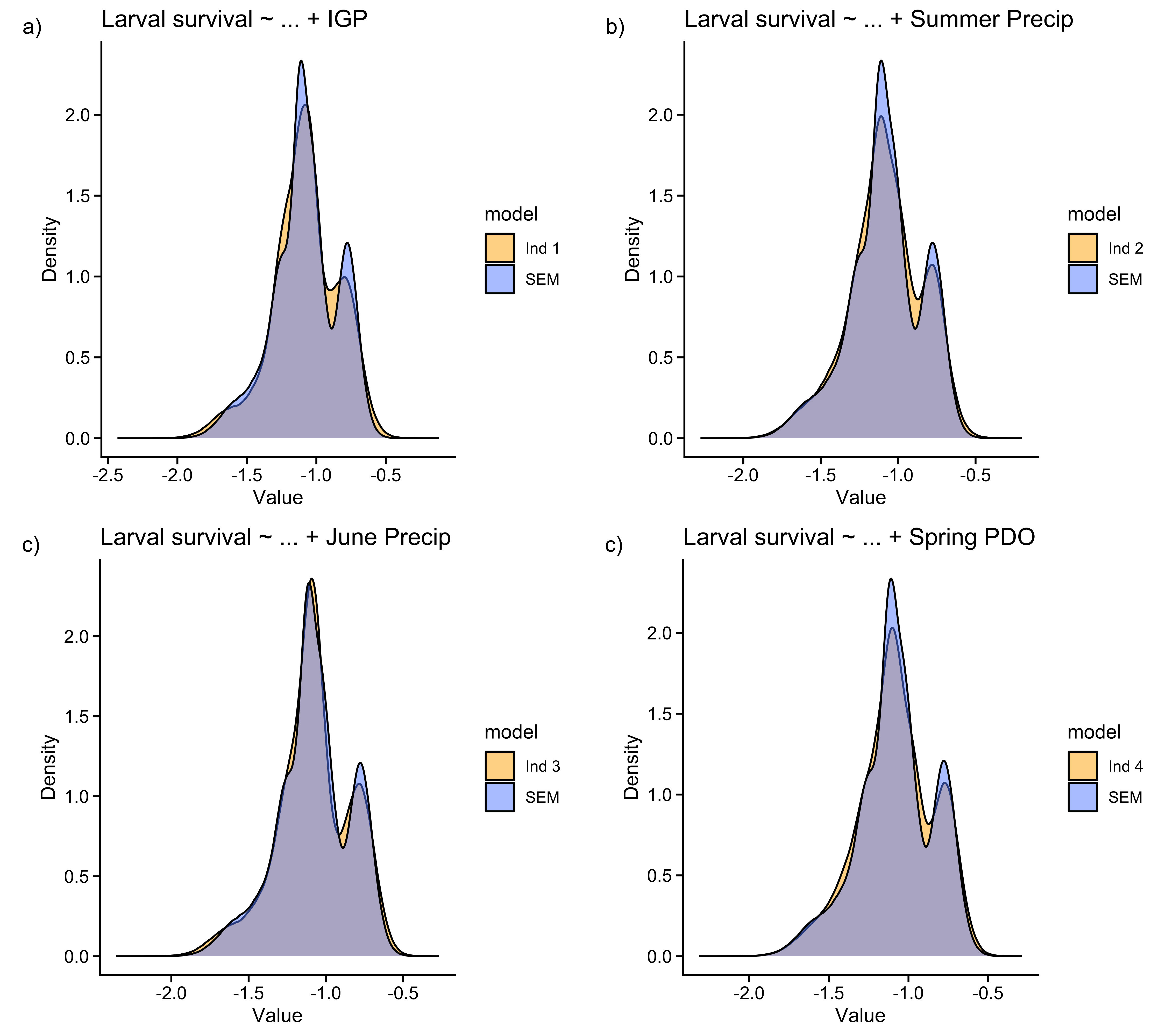

**Figure S3.** Density overlap plots of model predictions generated from the Bayesian structural equation model (SEM) for intraguild predation and its respective conditional independence tests. The SEM was defined by the following model structure shown in blue: Intraguild Predation ~ June Rainfall + Spring PDO. Yellow shows the addition of the tested independence claim to the model.

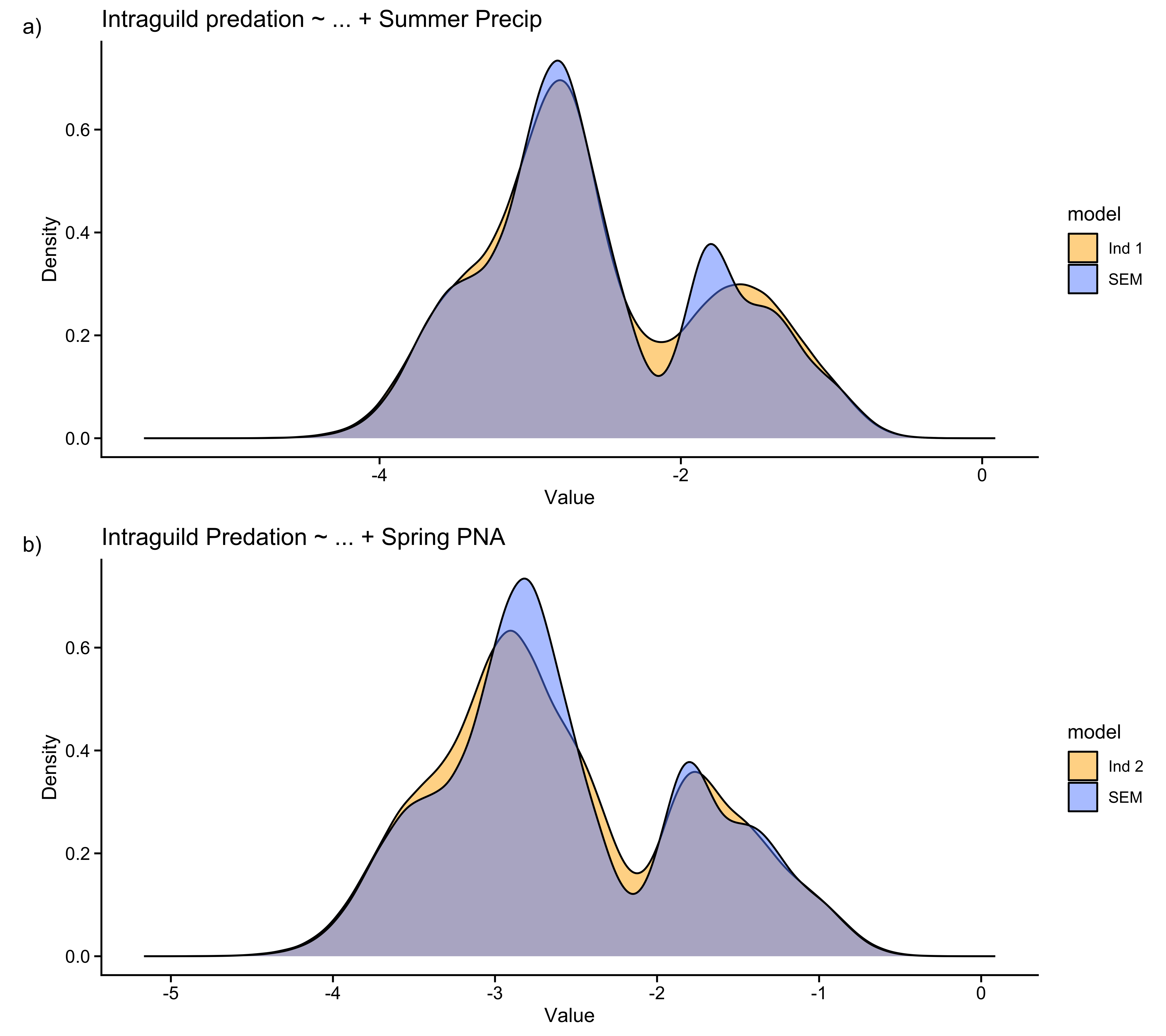

**Appendix B: Model Selection**

We began by reducing our candidate *a priori* predictor set for pre-diapause survival (Fig. S2) and intraguild predation (Fig. S3) by only retaining variables with a moderate effect (|ρ| > 0.5) size (Tredennick et al. 2021). We then further reduced the parameter space and eliminated multicollinearity between covariates by only keeping predictors whose variance inflation factors were all < 5 (Table S5; Dormann et al. 2013). To mitigate overfitting, we implemented a model selection approach based on regularization and Bayesian leave-one-out cross validation (loo-cv). Regularization was done using weakly informative shrinkage priors that retained parameters that only had strong effects (Table S6 & S7; McElreath, 2015; Lemoine, 2019). We used posterior predictive checks and a power-scaling sensitivity analysis to choose among priors that had similiar expected log predictive density (ELPD) values (Fig. S4 & S5; Table S7 & S8). To further reduce overfitting risk we applied a modified one-standard-error rule (m-OSE; Yates et al. 2022) and selection induced bias correction (McLatchie and Vehtari, 2024). The m-OSE reduces estimation uncertainty by favoring the least complex model whose adjusted error includes the estimate from the best-scoring model. We then estimated selection induced bias from the Bayesian loo-cv model comparisons and updated the ELPD values using a correction term (McLatchie and Vehtari, 2023). We then used the best model of pre-diapause survival and intraguild predation and tested for lag-1 temporal autocorrelation. After running the Bayesian structural equation model, we assessed goodness of fit in several ways. First, we ran a residual analysis where we plotted the relationship between the residuals and the fitted values for both model paths (Fig. S10). We then ran posterior predictive checks on the pre-diapause survival and intraguild predation model path (Fig. S11). We then examined several tests of conditional independence(Fig. S1 – S3).

**Figure S4.** Predictors of web survival retained after obtaining pairwise Pearson correlation coefficients (rho > 0.5).

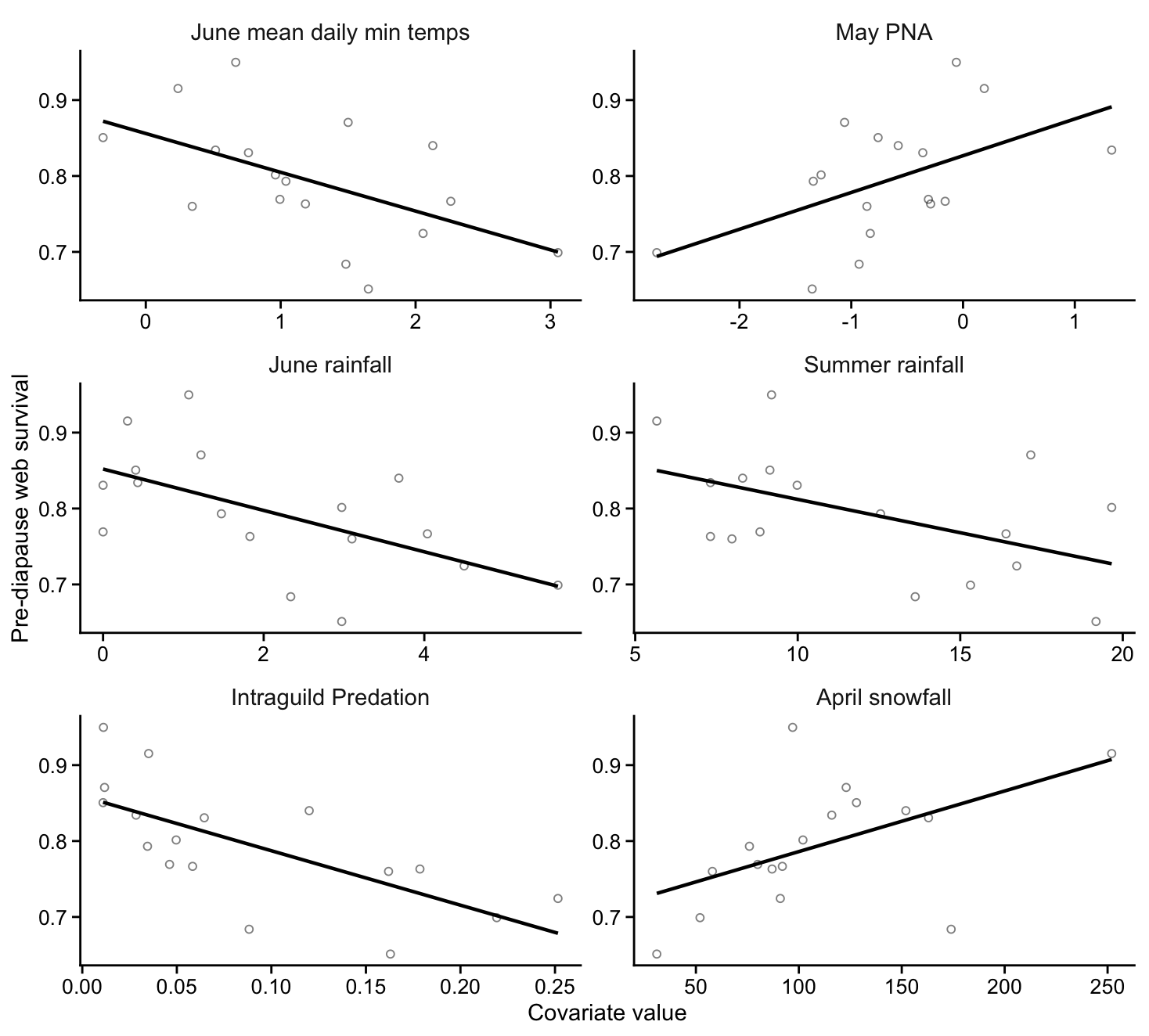

**Figure S5.** Predictors of larval survival retained after obtaining pairwise Pearson correlation coefficients (rho > 0.5).

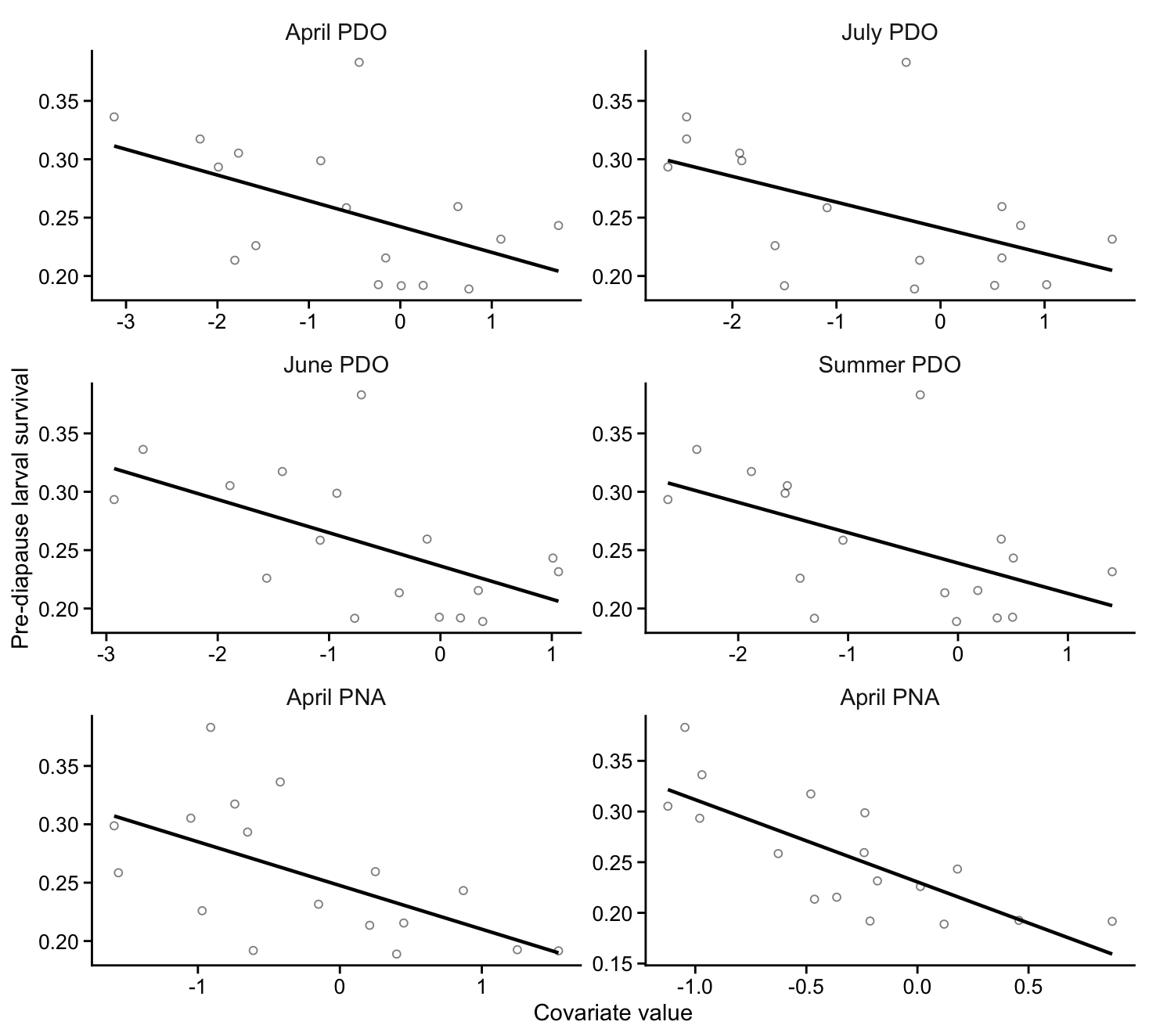

**Figure S6.** Predictors of intraguild predation retained after obtaining pairwise Pearson correlation coefficients (rho > 0.5).

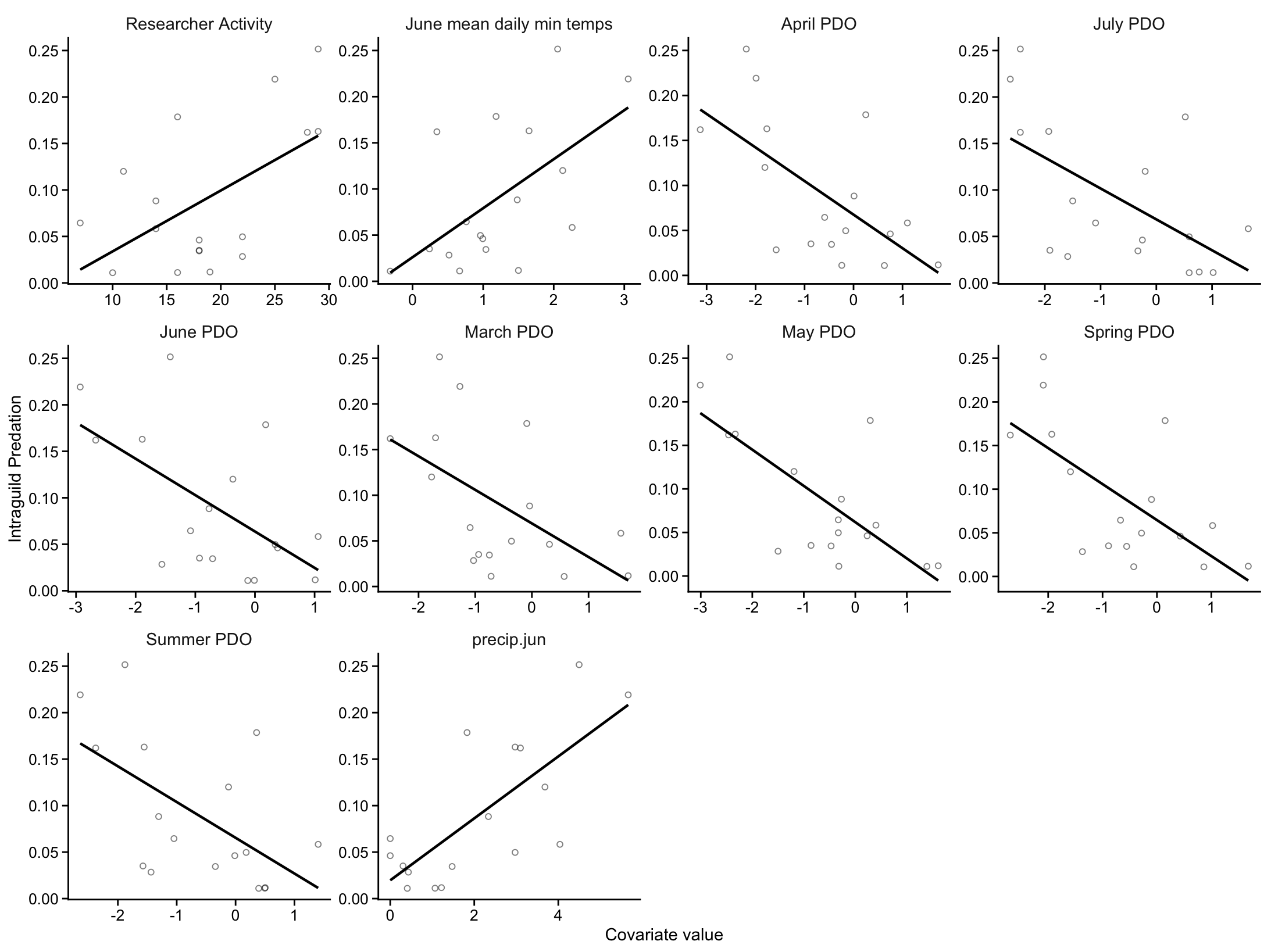

**Table S5.** Variance inflation factors among predictors of pre-diapause web & larval survival and intraguild predation.

| **Response** | **Predictor** | **VIF** |
| --- | --- | --- |
| Web survival | Intraguild Predation | 1.2 |
|  | Summer rainfall | 1.2 |
|  | April Snowfall | 1.4 |
| Larval Survival | Spring PNA | 2.0 |
|  | April PNA | 2.1 |
|  | Summer PDO | 1.3 |
| Intraguild Predation | Researcher Activity | 1.7 |
|  | June rainfall | 4.2 |
|  | Spring PDO | 1.7 |
|  | June daily minimum temps | 3.2 |

**Table S6.** Regularizing priors for web survival, larval survival, and intraguild predation ranked by expected log predictive density (ELPD), the effective number of parameters (P_loo), and the leave-one-out information criterion (LOOIC). Intercepts were fit with a prior based on the normal distribution, normal(0, 1). The Laplacian like prior was student_t(1, 0, 1), the normal prior was normal(0, 1), and the horseshoe prior was horseshoe(df = 1, par_ratio = 0.5). Pre-diapause survival ~ August mean daily maximum temps + Mean daily max temps from start of flight season to larval diapause + April PNA + Spring PNA. Intraguild Predation ~ Days of researcher activity + Spring PDO + June Rainfall. Convergence was assessed using 4 Markov chains that ran for 12000 iterations after discarding the first 2000 samples (total post-warmup draws = 40,000). For all models all Pareto k estimates were < 0.7.

| **Prior** | **ELPD** | **ELPD** | **Δ ELPD** | **P_loo** | **looic** |
| --- | --- | --- | --- | --- | --- |
|  | **Corrected** | **(SE)** | **(SE)** | **(SE)** | **(SE)** |
| Web survival |  |  |  |  |  |
| Horseshoe prior | -69.6 | -69.6 (12.2) |  | 11.2 (4.3) | 139.2 (24.4) |
| Laplacian prior | -70.1 | -69.9 (12.3) | -0.2 (1.4) | 12.2 (4.8) | 139.7 (24.5) |
| Normal prior | -70.2 | -69.9 (12.3) | -0.3 (1.4) | 12.3 (4.8) | 139.9 (24.5) |
| Larval survival |  |  |  |  |  |
| Laplacian prior | 28.7 | 28.7 (2.2) |  | 3.4 (0.9) | -57.5 (4.5) |
| Normal prior | 28.5 | 28.6 (2.3) | -0.1 (0.0) | 2.7 (0.5) | -57.3 (4.5) |
| Horseshoe prior | 28.4 | 28.6 (2.4) | -0.2 (0.6) | 3.0 (0.9) | -57.1 (4.7) |
| Intraguild Predation |  |  |  |  |  |
| Laplacian prior | 28.5 | 28.5 (4.3) |  | 3.9 (1.2) | -57.0 (8.6) |
| Normal prior | 28.4 | 28.5 (4.3) | -0.0 (0.1) | 4.0 (1.3) | -56.9 (8.7) |
| Horsehoe prior | 26.5 | 26.5 (3.5) | -2.0 (1.1) | 4.2 (7.1) | -53.0 (7.1) |

**Figure S7.** Kernel density estimate of the observed values (dark line) for web survival when fit using (a) laplace like, (b) normal, and (c) horseshoe priors with density estimates for 500 simulated values (light blue lines) drawn from the posterior predictive distribution.

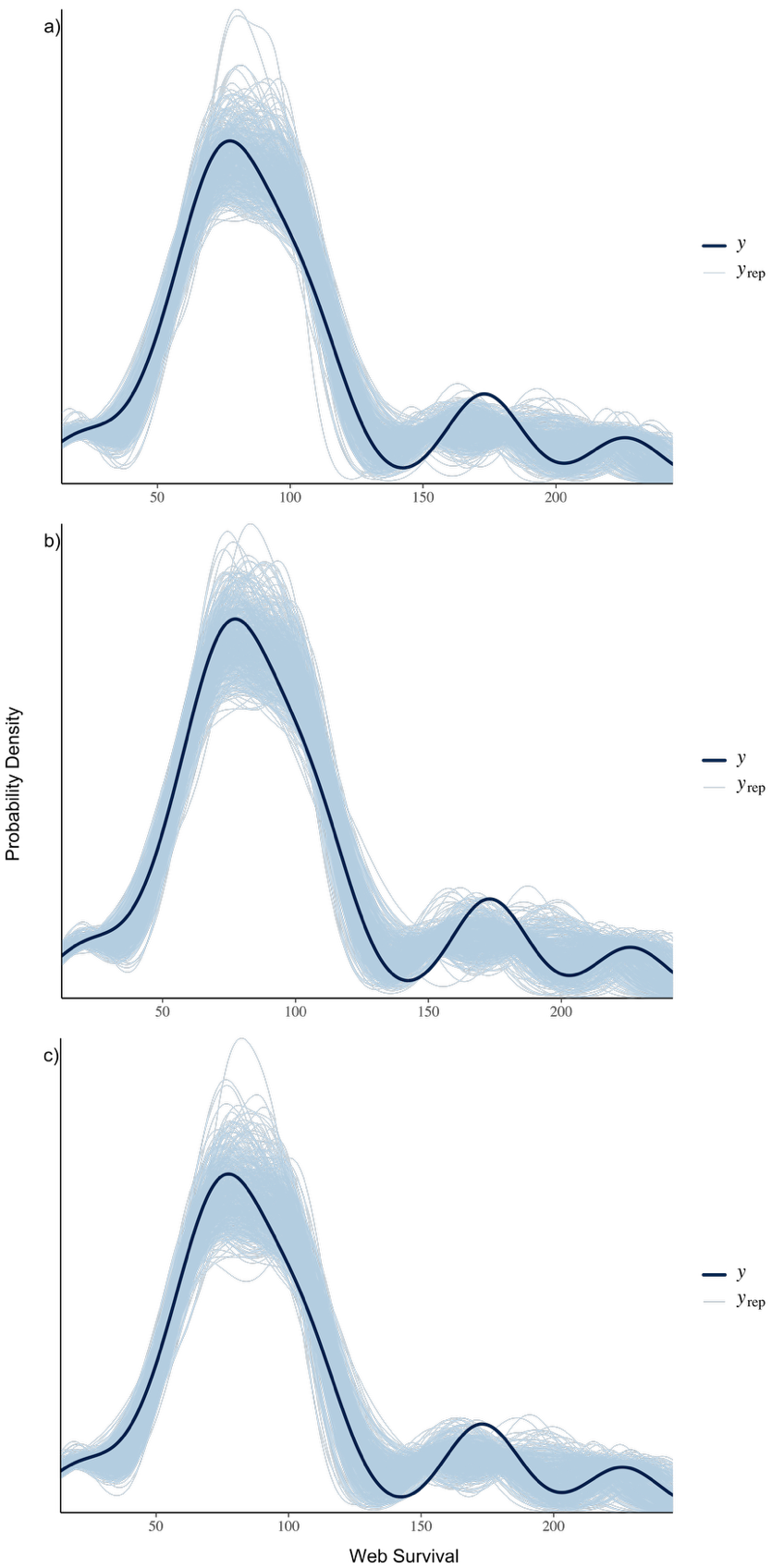

**Table S7.** Results from the power-scaling sensitivity analysis used to assess priors describing the web survival Bayesian structural equation model path. The prior for beta coefficients was defined as student_t(1,0,1); the intercept prior was defined as normal(0,1).

| **Predictor** | **Prior** | **Likelihood** | **Diagnosis** |
| --- | --- | --- | --- |
| b_Intercept | 0.014 | 0.074 |  |
| Intraguild Predation | 0.012 | 0.081 |  |
| Summer rainfall | 0.011 | 0.075 |  |
| April snowfall | 0.007 | 0.079 |  |
| Intercept | 0.014 | 0.074 |  |
| Prior intercept | 0.000 | 0.002 |  |
| Prior b | 0.000 | 0.002 |  |

**Figure S8.** Kernel density estimate of the observed values (dark line) for larval survival when fit using (a) laplace like, (b) normal, and (c) horseshoe priors with density estimates for 500 simulated values (light blue lines) drawn from the posterior predictive distribution.

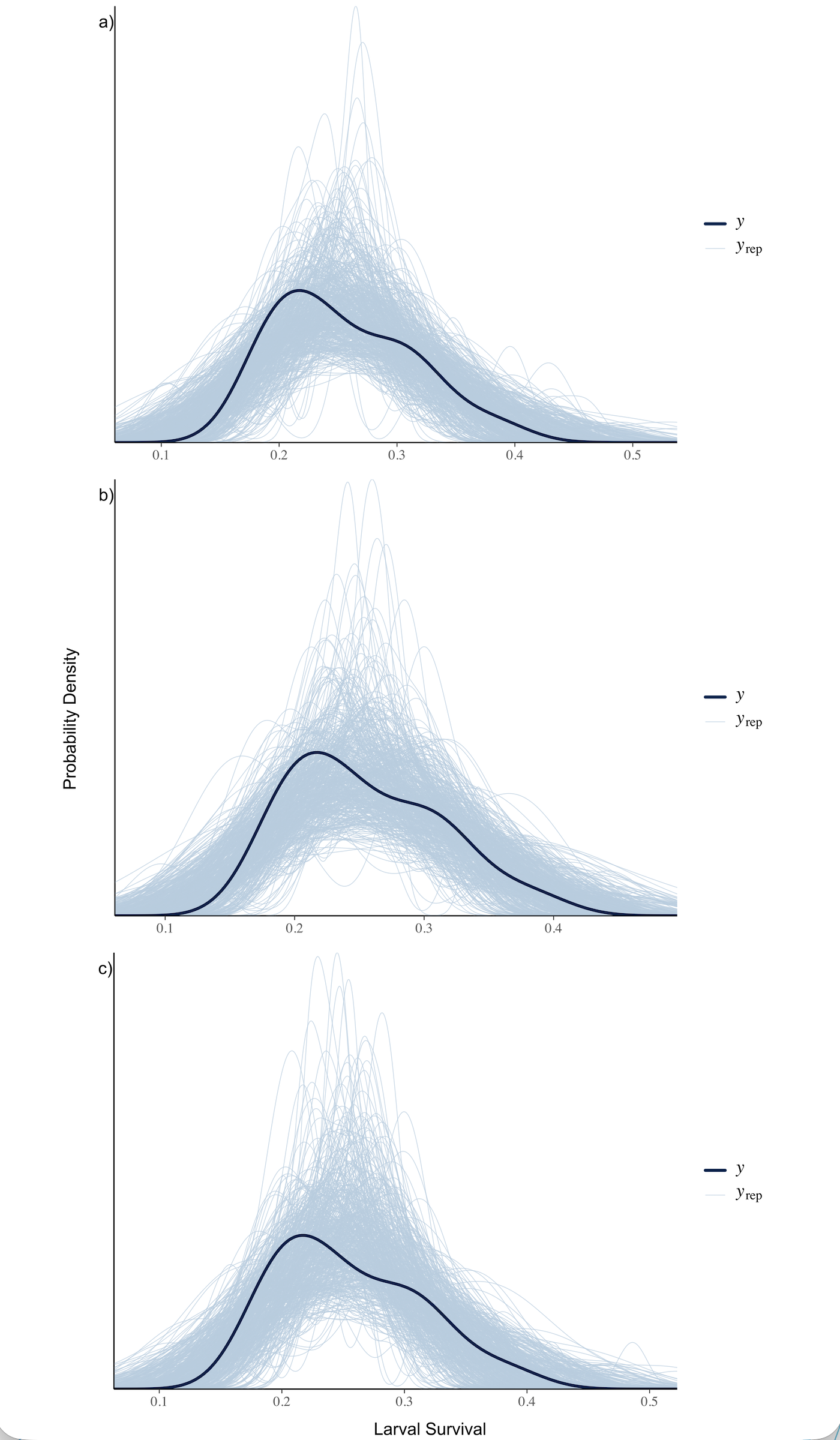

**Table S8.** Results from the power-scaling sensitivity analysis used to assess priors describing the larval survival Bayesian structural equation model path. The prior for beta coefficients was defined as student_t(1,0,1); the intercept prior was defined as normal(0,1). The initial prior for phi was gamma(0.01, 0.01). Phi was then modified sequentially until there no longer remained any diagnostic warnings; the final phi prior was gamma(2, 0.005).

| **Prior set** | **Predictor** | **Prior** | **Likelihood** | **Diagnosis** |
| --- | --- | --- | --- | --- |
| Initial | b_Intercept | 0.034 | 0.114 |  |
|  | Spring PNA | 0.026 | 0.107 |  |
|  | April PNA | 0.019 | 0.099 |  |
|  | Summer PDO | 0.018 | 0.100 |  |
|  | Phi | 0.168 | 0.197 | Potential prior-data conflict |
|  | Intercept | 0.034 | 0.114 |  |
|  | Prior intercept | 0.001 | 0.003 |  |
|  | Prior b | 0.005 | 0.003 |  |
|  | Prior phi | 0.036 | 0.042 |  |
| Final | b_Intercept | 0.008 | 0.079 |  |
|  | Spring PNA | 0.014 | 0.080 |  |
|  | April PNA | 0.010 | 0.084 |  |
|  | Summer PDO | 0.005 | 0.085 |  |
|  | Phi | 0.013 | 0.078 |  |
|  | Intercept | 0.008 | 0.079 |  |
|  | Prior intercept | 0.000 | 0.003 |  |
|  | Prior b | 0.000 | 0.002 |  |
|  | Prior phi | 0.000 | 0.007 |  |

**Figure S9.** Kernel density estimate of the observed values (dark line) for intraguild predation when fit using (a) laplace, (b) normal, and (c) horseshoe priors with density estimates for 500 simulated values (light blue lines) drawn from the posterior predictive distribution.

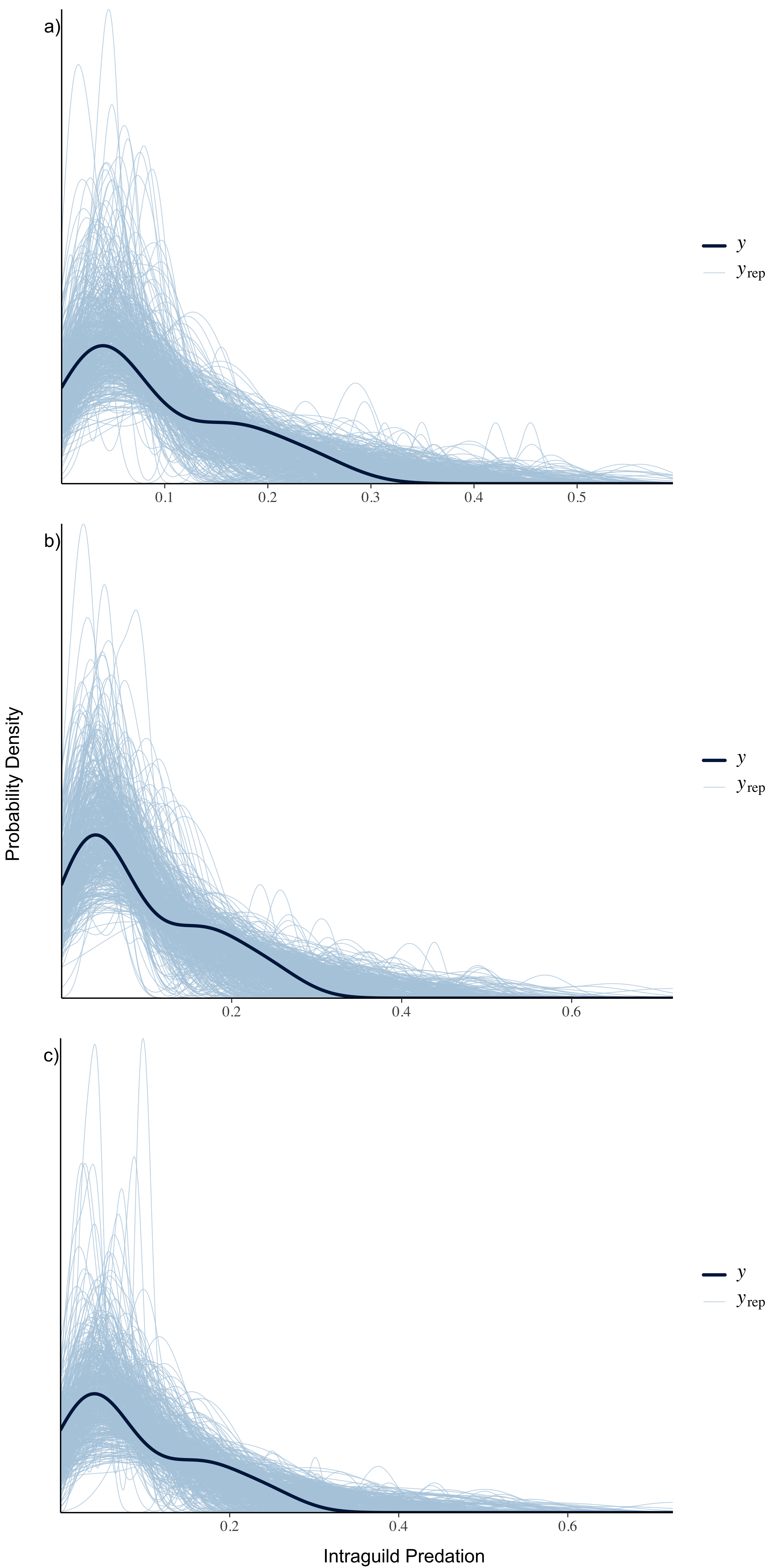

**Table S9.** Results from the power-scaling sensitivity analysis used to assess priors describing the intraguild predation Bayesian structural equation model path. The initial prior for beta coefficients was defined as student_t(1,0,1); the intercept prior was defined as normal(0,1); and the prior for phi was gamma(0.01, 0.01). All priors were then modified sequentially until there no longer remained any diagnostic warnings; the final beta coefficient prior was student_t(2,0,2); the intercept was normal(0,2); and phi was gamma(2, 0.005).

| **Prior set** | **Predictor** | **Prior** | **Likelihood** | **Diagnosis** |
| --- | --- | --- | --- | --- |
| Initial | b_Intercept | 0.129 | 0.191 | Potential prior-data conflict |
|  | Researcher Activity | 0.008 | 0.078 |  |
|  | June Rainfall | 0.035 | 0.070 |  |
|  | Spring PDO | 0.053 | 0.109 | Potential prior-data conflict |
|  | June daily min | 0.026 | 0.064 |  |
|  | Phi | 0.154 | 0.169 | Potential prior-data conflict |
|  | Intercept | 0.129 | 0.191 | Potential prior-data conflict |
|  | Prior intercept | 0.002 | 0.004 |  |
|  | Prior b | 0.006 | 0.011 |  |
|  | Prior phi | 0.024 | 0.046 |  |
| Final | b_Intercept | 0.011 | 0.085 |  |
|  | Researcher Activity | 0.011 | 0.091 |  |
|  | June Rainfall | 0.019 | 0.076 |  |
|  | Spring PDO | 0.017 | 0.078 |  |
|  | June daily min | 0.017 | 0.075 |  |
|  | Phi | 0.043 | 0.083 |  |
|  | Intercept | 0.011 | 0.085 |  |
|  | Prior intercept | 0.001 | 0.003 |  |
|  | Prior b | 0.001 | 0.003 |  |
|  | Prior phi | 0.001 | 0.011 |  |

**Figure S10.** Plots of the relationship between residuals and fitted values for (a) pre-diapause survival and (b) intraguild predation.

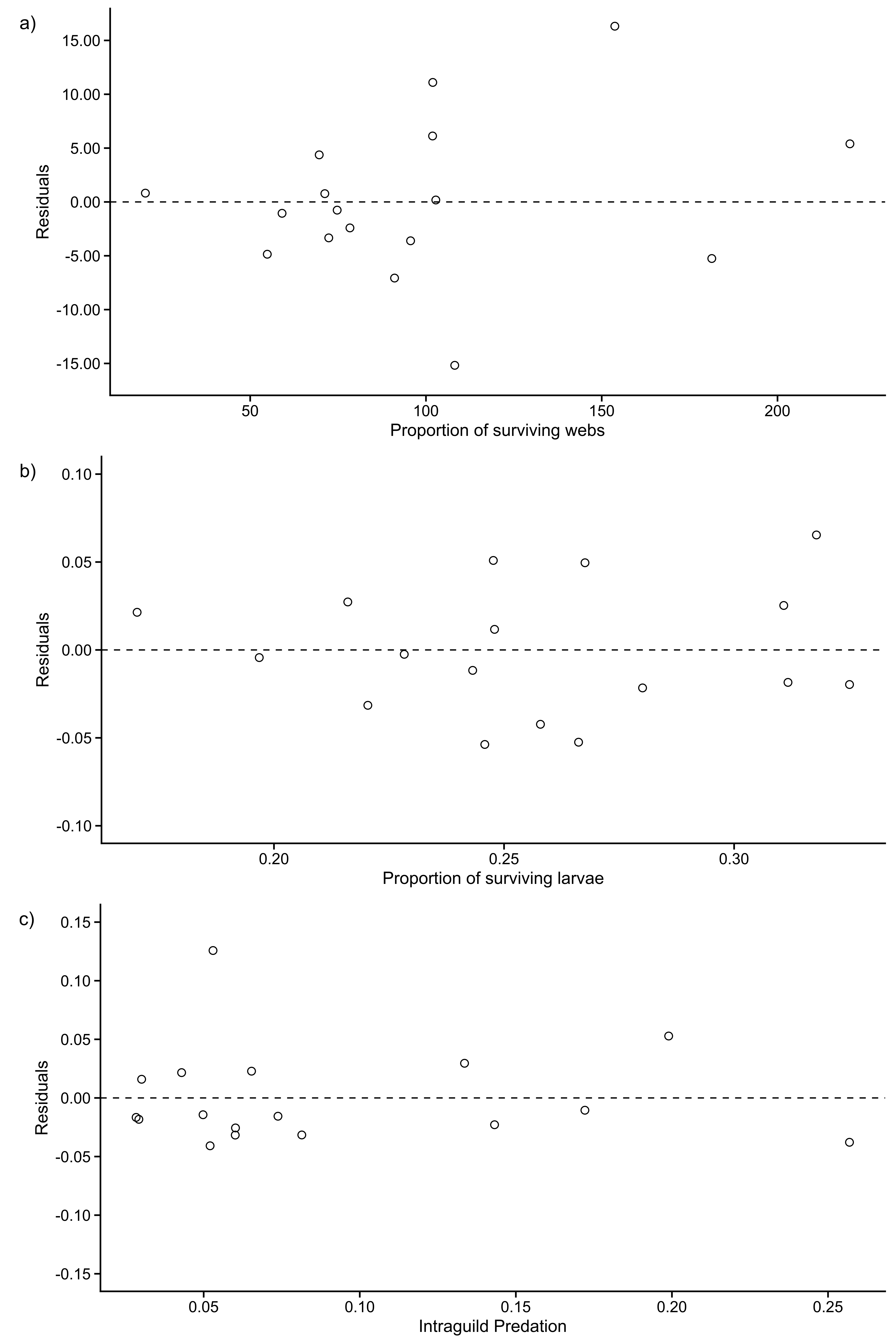

**Figure S11.** Kernel density estimate of the observed values (dark line) for (a) web survival, (b) larval survival, and (c) intraguild predation with density estimates for 500 simulated values drawn from the posterior predictive distribution of the final Bayesian structural equation model.

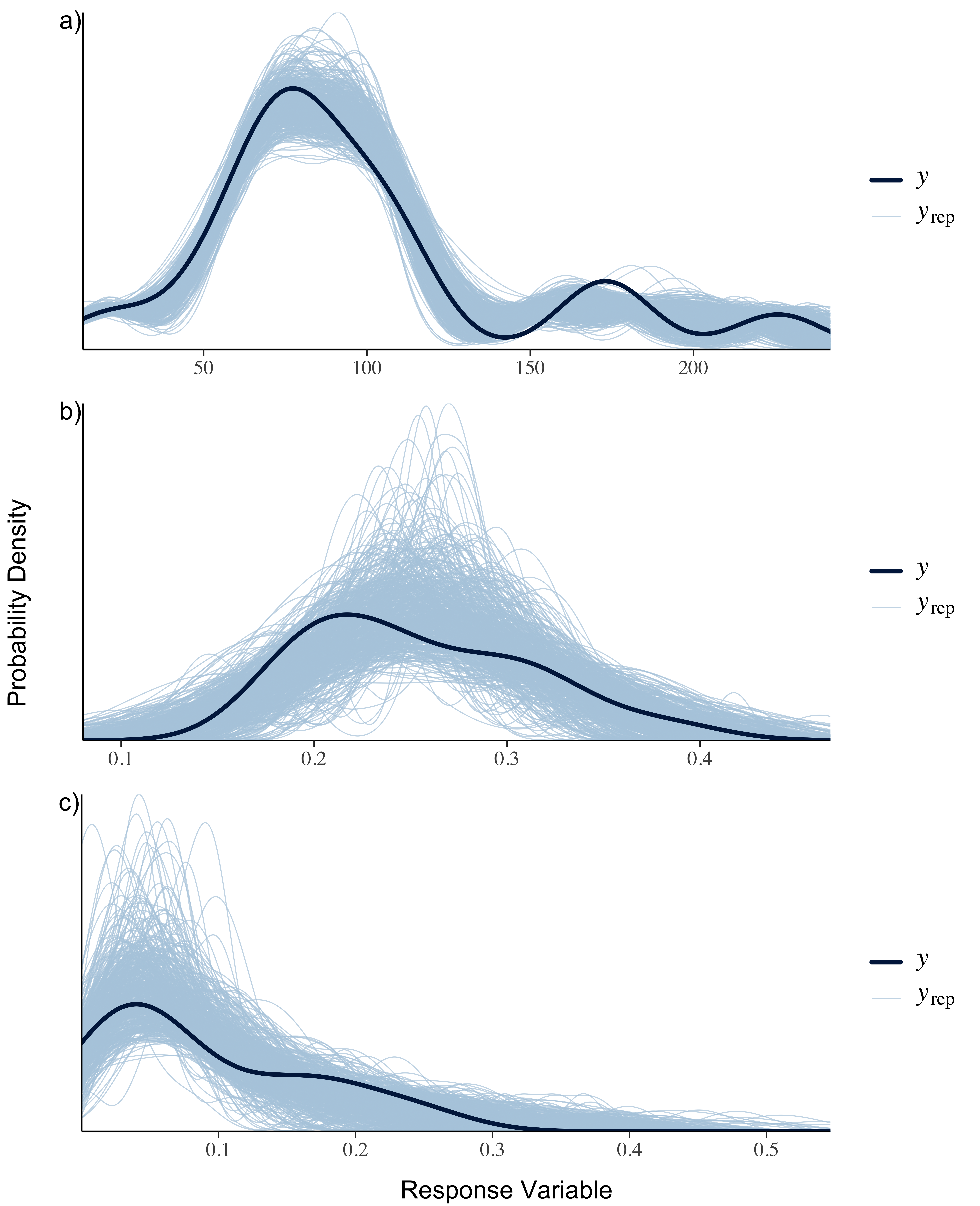
